# Mitochondrial phenotypes in purified human immune cell subtypes and cell mixtures

**DOI:** 10.1101/2020.10.16.342923

**Authors:** Shannon Rausser, Caroline Trumpff, Marlon A McGill, Alex Junker, Wei Wang, Siu-hong Ho, Anika Mitchell, Kalpita R Karan, Catherine Monk, Suzanne C. Segerstrom, Rebecca G. Reed, Martin Picard

## Abstract

Using a high-throughput mitochondrial phenotyping platform to quantify multiple mitochondrial features among molecularly-defined immune cell subtypes, we quantify the natural variation in citrate synthase, mitochondrial DNA copy number (mtDNAcn), and respiratory chain enzymatic activities in human neutrophils, monocytes, B cells, and naïve and memory T lymphocyte subtypes. In mixed peripheral blood mononuclear cells (PBMCs) from the same individuals, we show to what extent mitochondrial measures are confounded by both cell type distributions and contaminating platelets. Cell subtype-specific measures among women and men spanning 4 decades of life indicate potential age- and sex-related differences, including an age-related elevation in mtDNAcn, which are masked or blunted in mixed PBMCs. Finally, a proof-of-concept, repeated-measures study in a single individual validates cell type differences and also reveals week-to-week changes in mitochondrial activities. Larger studies are required to validate and mechanistically extend these findings. These mitochondrial phenotyping data build upon established immunometabolic differences among leukocyte sub-populations, and provide foundational quantitative knowledge to develop interpretable blood-based assays of mitochondrial health.

## Introduction

Mitochondria are the most studied organelle across the biomedical sciences (Picard, Wallace, & Burelle, 2016). The growing focus on mitochondria is motivated by evidence positioning mitochondrial (dys)function as a driver of disease risk and aging (Jang, Blum, Liu, & Finkel, 2018; Picard et al., 2016; Wallace, 2015), and as a mediator of brain-body processes that shape health and disease trajectories across the lifespan (Picard, Trumpff, & Burelle, 2019). Addressing emerging biomedical research questions around the role of mitochondria on human health requires tractable quantitative biomarkers of mitochondrial content (the amount or mass of mitochondria per cell) and function (energy production capacity) that can be deployed in accessible human tissues, such as peripheral blood leukocytes. To develop such biomarkers, we need to establish standard effect sizes of mitochondrial variation – between immune cell subtypes and over time – as well as a quantitative handle on potential technical confounds and covariates such as age, sex, and known biomarkers.

Although some cell-specific assays can interrogate immune cells mitochondrial function with a reasonable degree of cell specificity (Chacko et al., 2013), more frequent approaches in the literature use peripheral blood mononuclear cells (PBMCs) (Dixon et al., 2019; Ehinger, Morota, Hansson, Paul, & Elmér, 2016; Karabatsiakis et al., 2014; Picard et al., 2018; Tyrrell et al., 2015; Weiss et al., 2015). These approaches largely assume that the immunometabolic properties of different immune cells have a negligible influence on mitochondrial measurements. However, there are marked differences in the metabolic properties of different immune cell subtypes well known to immunologists. For example, lymphocytes and monocytes significantly differ in their respiratory properties and mitochondrial respiratory chain (RC) protein abundance (Chacko et al., 2013; Kramer, Ravi, Chacko, Johnson, & Darley-Usmar, 2014; Maianski et al., 2004; Pyle et al., 2010). In various leukocyte subtypes, these divergent immunometabolic properties even contribute to determine the acquisition of specialized cellular characteristics (Pearce, Poffenberger, Chang, & Jones, 2013). The activation, proliferation, and differentiation of monocytes (Nomura et al., 2016) and T cells (Michalek et al., 2011) into specific effector cells require distinct metabolic profiles and cannot proceed without the proper metabolic states. Likewise, naïve and memory T lymphocytes differ in their reliance on mitochondrial oxidative phosphorylation (OxPhos) involving the RC enzymes (K. Brand, 1985; Jones et al., 2019; Ron-Harel et al., 2019), and harbor differences in protein composition and mitochondrial content within the cytoplasm (Bektas et al., 2019). Thus, the immune system offers a well-defined landscape of metabolic profiles which, if properly mapped, can potentially serve as biomarkers.

The composition of peripheral blood leukocytes in the human circulation is influenced by several factors. Immune cell subtypes are normally mobilized from lymphoid organs into circulation in a diurnal fashion and with acute stress (Ackermann et al., 2012; Beis et al., 2018; Dhabhar, Malarkey, Neri, & McEwen, 2012; Dhabhar, Miller, Stein, McEwen, & Spencer, 1994). The abundance of circulating immune cell subtypes also vary extensively between individuals, partially attributable to both individual-level (e.g., sex and age) and environmental factors (Patin et al., 2018). As a result, sampling whole blood or mixed PBMCs from different individuals reflect different cell populations that may not be directly comparable.

Furthermore, frequently used Ficoll-isolated PBMCs are naturally contaminated with (sticky) platelets (Butler et al., 2007). Platelets contain mitochondria and mtDNA but no nuclear genome to use as reference for mtDNA copy number (mtDNAcn) measurements (Hurtado-Roca et al., 2016), representing a major source of bias to mitochondrial studies in PBMCs or other undefined cell mixtures (Banas, Kost, & Goebel, 2004; Shim, Arshad, Gadawska, Côté, & Hsieh, 2020; Urata, Koga-Wada, Kayamori, & Kang, 2008). But the extent to which platelet contamination influences specific mitochondrial content and RC capacity features in PBMCs has not been quantitatively defined.

Another significant gap in knowledge relates to the natural dynamic variation in mitochondrial content and function over time. Mitochondria dynamically recalibrate their form and functions in response to mental stress (Picard & McEwen, 2018) and exercise (Gan, Fu, Kelly, & Vega, 2018). Mitochondria also contain receptors that enable their functions to be tuned by humoral metabolic and endocrine inputs (Bénard et al., 2012; Du et al., 2009). Thus, cell-specific mitochondrial features could vary over time. To develop valid blood-based mitochondrial markers, we therefore need to determine whether leukocyte mitochondria are stable *trait*-like properties of each person, or *state*-like properties possibly varying in response to metabolic or endocrine factors.

To address these questions, we used a high-throughput mitochondrial phenotyping platform on immunologically-defined immune cell subtypes, in parallel with PBMCs, to quantify cell subtype-specific mitochondrial phenotypes in a small, diverse cohort of healthy adults. First, we establish the extent to which cell type composition and platelet contamination influence PBMC-based mitochondrial measures. We then systematically map the mitochondrial properties of different immune cell subtypes and validate the existence of stable mitochondrial phenotypes in an intensive repeated-measures design within the same individual, which also begins to reveal a surprising degree of intra-individual variation over time. Collectively, these data confirm and quantify the biological limitations of PBMCs to profile human mitochondria, introduce the concept of multivariate mitotypes, and define unique cell-specific mitochondrial features in circulating human leukocytes in relation to age, sex, and biomarkers that can guide future studies. Together, these data represent a resource to design cell-specific immune mitochondrial phenotyping strategies.

## Results

### Cell subtype distributions by age and sex

We performed mitochondrial profiling on molecularly-defined subtypes of immune cell populations in parallel with PBMCs in twenty-one participants (11 women, 10 men) distributed across 4 decades of life (ages 20-59, 4-8 participants per decade across both sexes). From each participant, 100ml of blood was collected; total leukocytes were then labeled with two cell surface marker cocktails, counted, and isolated by fluorescence-activated cell sorting (FACS, see Methods for details), frozen, and subsequently processed as a single batch on our mitochondrial phenotyping platform adapted from (Picard et al., 2018) (Figure 1a). In parallel, a complete blood count (CBC) for the major leukocyte populations in whole blood, standard blood chemistry, and a metabolic and endocrine panel were assessed (Figure 1b).

**Figure 1.**
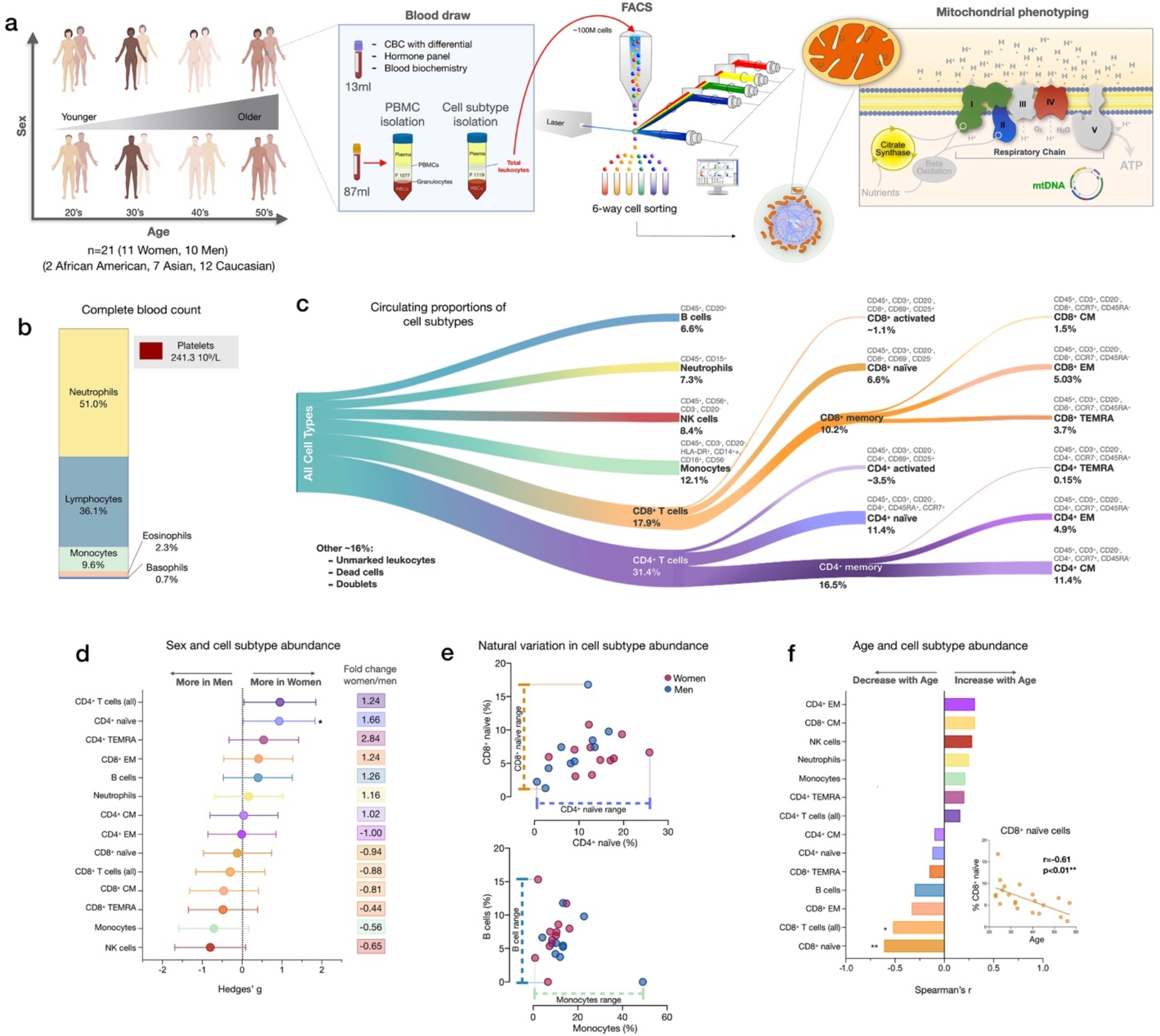
Immune cell subtype distribution in adult women and men. (**a**) Overview of participant demographics, blood collection, processing, and analysis pipeline. Total leukocytes were isolated using Ficoll 1119 and PBMCs were isolated on Ficoll 1077. (*right*) The five mitochondrial features analyzed on the mitochondrial phenotyping platform are colored. Mitochondrial phenotyping platform schematic adapted from Figure 1A (Picard et al., 2018). (**b**) Stacked histogram showing the leukocytes distribution derived from the complete blood count (CBC) of major cell types. (**c**) Diagram illustrating the proportion of circulating immune cell subtypes (% of all detected cells) quantified by flow cytometry from total peripheral blood leukocytes. Cell surface markers and subtype definitions are detailed in Supplementary file 1. (**d**) Effect sizes for cell subtype distribution differences between women (n=11) and men (n=10). P-values from non-parametric Mann-Whitney T test. Error bars reflect the 95% confidence interval (C.I.) on the effect size, and the fold change comparing raw counts between women and men is shown on the right. (**e**) Example distributions of cell type proportions in women and men illustrating the range of CD4^+^ and CD8^+^ naïve cells, B cells, and monocytes, highlighting the natural variation among our cohort. Each datapoint reflects a different individual. (**f**) Spearman’s r correlation between age and cell types proportion. n=21, p<0.05*, p<0.01**.

We first quantified the abundance of specific cell subtypes based on cell surface marker combinations (Figure 1c, Figure 1-figure supplements 1-2, and Supplementary file 1). Men had 66% fewer CD4^+^ naïve T cells than women (p<0.05) and tended to have on average 35-44% more NK cells and monocytes (Figure 1d). These differences were characterized by moderate to large standardized effect sizes (Hedge’s g=0.71-0.93), consistent with recent findings (Márquez et al., 2020). Between individuals of the same sex, the circulating proportions of various cell subtypes (e.g., B cells range: <0.01-15.3%, see Supplementary file 2) varied by up to an order of magnitude (i.e., 10-fold) (Figure 1e).

In relation to age, as expected (Patin et al., 2018), CD8^+^ naïve T cell abundance was lower in older individuals (p<0.01). Compared to young adults in their 20’s, middle-aged individuals in their 50’s had on average ~63% fewer CD8^+^ naïve T cells (Figure 1f). In contrast, effector memory CD4^+^ (CD4^+^ EM) and central memory CD8^+^ (CD8^+^ CM) cell abundance tended to increase with age (positive correlation, r=0.31 for both), an overall picture consistent with immunological aging (Márquez et al., 2020; Nikolich-Žugich, 2014; Patin et al., 2018).

CBC-derived cell proportions also showed that men had on average 28% more monocytes than women (Figure 1-figure supplement 3), consistent with our FACS results. Conversely, women had on average 20% more platelets than men. Platelet abundance also tended to decrease with age, a point discussed later.

### Circulating cell composition influence PBMCs mitochondrial phenotypes

We next examined how much the abundance of various circulating immune cell subtypes correlated with individual mitochondrial metrics in PBMCs. Our analysis focused on two key aspects of mitochondrial biology: i) *mitochondrial content*, indexed by citrate synthase (CS) activity, a Kreb’s cycle enzyme used as a marker of mitochondrial volume density (Larsen et al., 2012), and mtDNAcn, reflecting the number of mtDNA copies per cell; and ii) *RC function* measured by complex I (CI), complex II (CII) and complex IV (CIV) enzymatic activities, which reflect the capacity for electron transport and respiratory capacity and serve here as a proxy for maximal RC capacity. Furthermore, by adding the three mean-centered features of RC function together as a numerator (CI+CII+CIV), and dividing this by the combination of content features (CS+mtDNAcn), we obtained an index reflecting *RC capacity on a per-mitochondrion basis*, known as the mitochondrial health index (MHI) adapted from previous work (Picard et al., 2018).

As expected, the abundance of multiple circulating cells was correlated with PBMCs mitochondrial features (Figure 2). Notably, the correlation between circulating B cell abundance and PBMCs CS activity was r=0.78 (p<0.0001), meaning that the proportion of shared variance (r^2^) between both variables is 61% (i.e., B cell abundance explains 61% of the variance in PBMCs CS). Similarly, the correlations between B cell abundance and PBMCs mtDNAcn, CI, CII activities were 0.52-0.67 (27-45% of shared variance, ps<0.05-01). The circulating proportions of other cell types accounted for more modest portions (r^2^<14%) of the variance in PBMCs, although the higher abundance of memory cells tended to be negatively associated with PBMC RC enzymatic activities.

**Figure 2.**
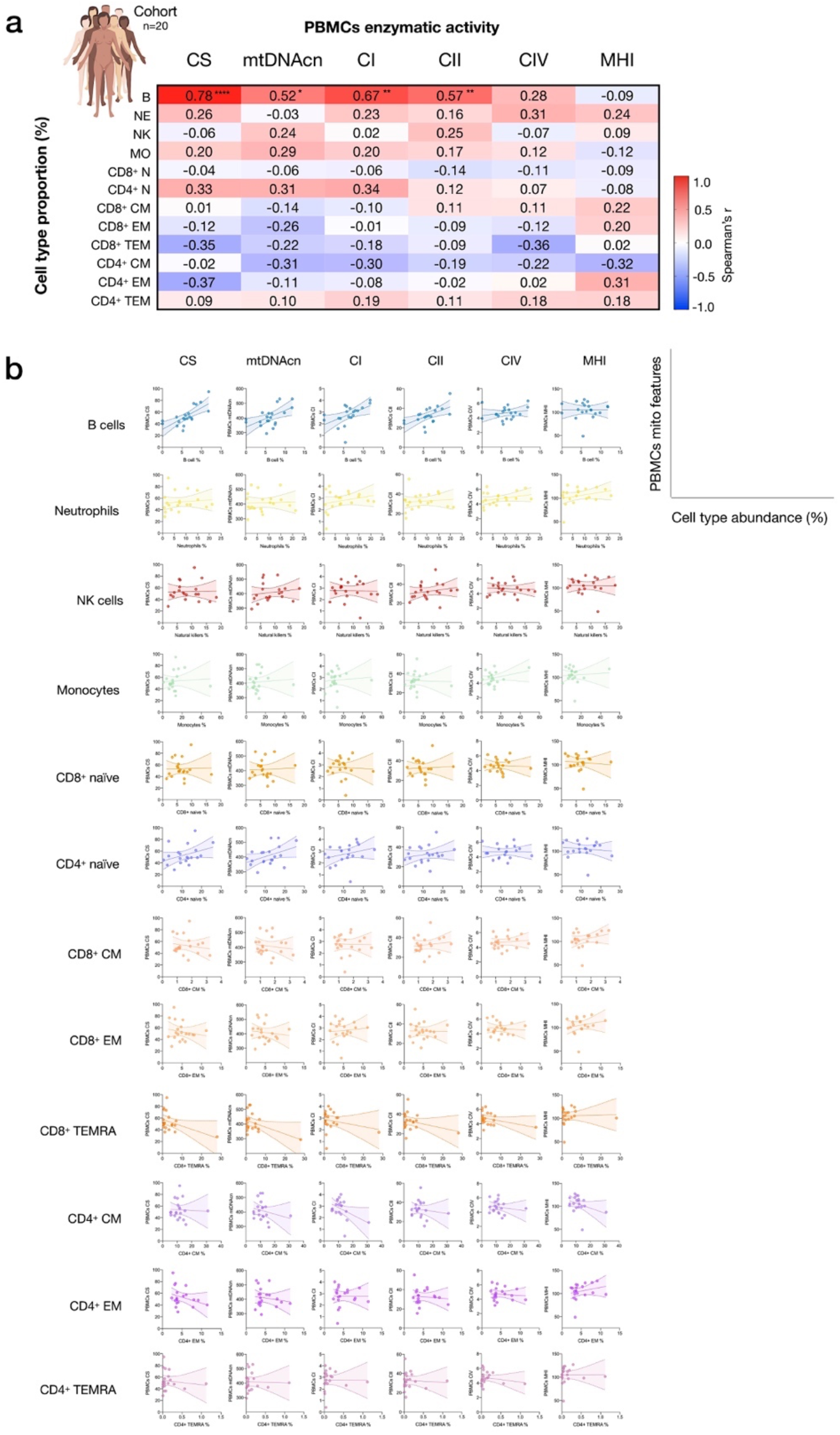
Influence of cell subtypes on mitochondrial features in total PBMCs. (**a**) Pairwise correlations (Spearman’s r) between cell subtype proportions obtained from cell sorting with mitochondrial features measured in PBMCs for the cohort (n=20, 1 participant with missing PBMC-measured data). Aggregate correlations are shown as a heatmap (*top*) and (**b**) individual scatterplots (*bottom*).

Based on CBC-derived cell proportions, the abundance of eosinophils and neutrophils was positively correlated with most PBMC mitochondrial content and activity features (Figure 2-figure supplement 1). Because PBMCs do not contain granulocytes, these correlations may reflect the independent effect of a humoral factor on cell mobilization and mitochondrial function. Together, these data confirmed that mitochondrial features assessed in PBMCs in part reflect the proportions of some but not all circulating cell subtypes, quantitatively documenting how cell type distribution may confound the measurements of mitochondrial function in PBMCs.

### Platelets influence PBMCs mitochondrial phenotypes

Given that platelets easily form platelet-leukocyte aggregates (Butler et al., 2007) (Figure 3a), to partly resolve the origin of the discrepancies between isolated cell subtypes and PBMCs noted above, we directly quantified the contribution of platelets to total mitochondrial content and activity features in PBMCs. We note that the PBMCs in our experiments were carefully prepared with two passive platelet depletion steps (low speed centrifugations, see Appendix 1 for details), which should have already produced “clean” PBMCs.

**Figure 3.**
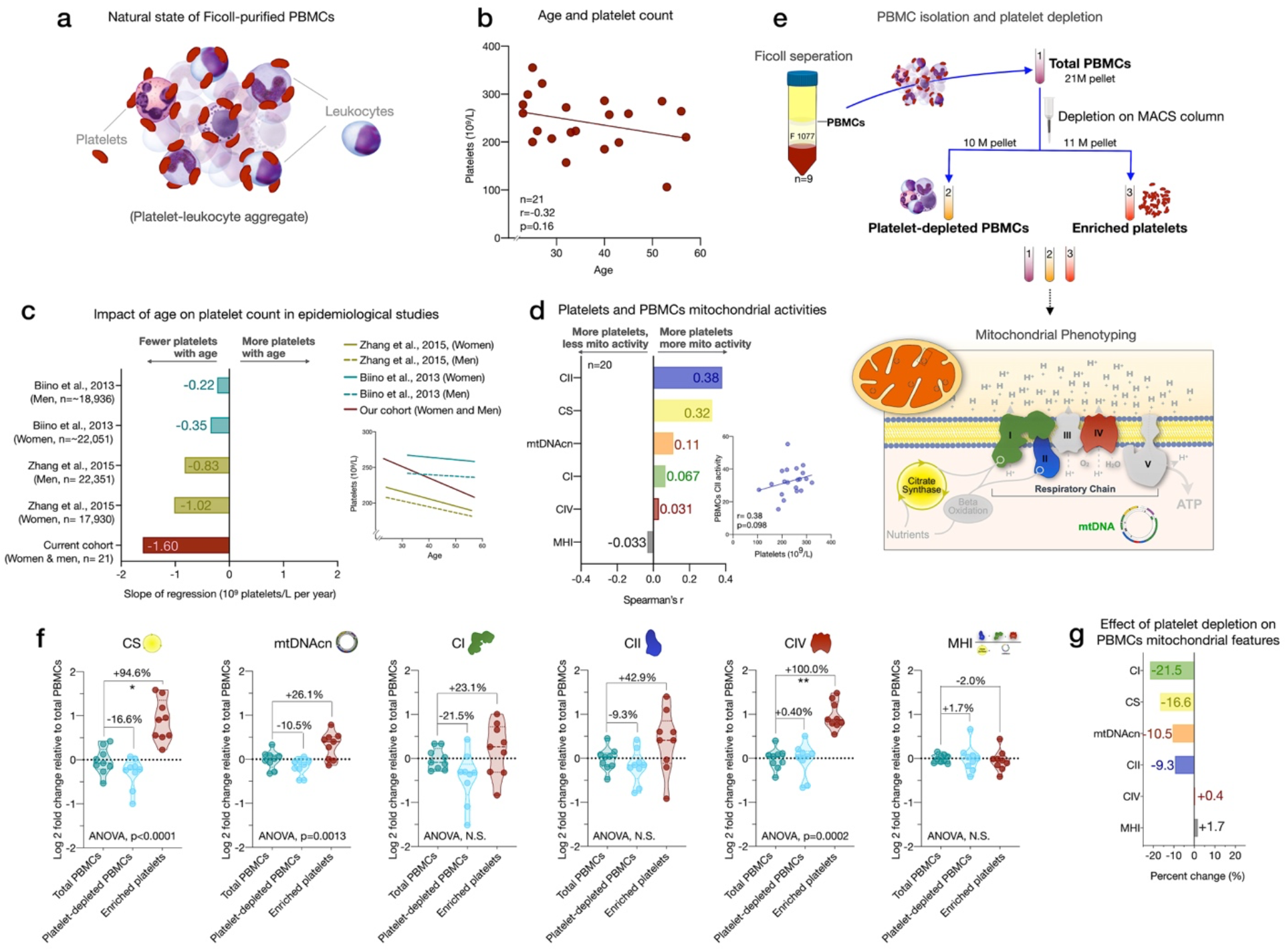
Influence of platelet contamination on mitochondrial features in total PBMCs. (**a**) Schematic of the natural state of Ficoll-isolated PBMCs associated with contaminating platelets. (**b**) Association of age and circulating platelet abundance (i.e., count) in our cohort (Spearman’s r). (**c**) Change in platelet abundance as a function of age. The magnitude of the association (slope of the regression: 109 platelets/L per year) from two large epidemiological studies and our cohort. The inset shows the actual regressions (n=21 to 22,351). (**d**) Effect sizes of the association between platelet count and PBMC mitochondrial features in our cohort (n=20). (**e**) Overview of the experimental PBMC platelet depletion study, yielding three different samples subjected to mitochondrial phenotyping. Mitochondrial phenotyping platform schematic adapted from Figure 1A (Picard et al., 2018). (**f**) Fold change in mitochondrial parameters between i) platelet-depleted PBMCs, and ii) enriched platelets (with contaminating PBMCs), relative to iii) total PBMCs. P-values from One-Way non-parametric ANOVA Friedman test, post-hoc Dunn’s multiple comparisons relative to total PBMCs. (**g**) Percent change of platelet-depleted PBMCs mitochondrial features from total PBMCs. n=9, p<0.05*, p<0.01**, p<0.001***, p<0.0001****.

We first asked if the abundance of platelets from the CBC data in the cohort varies by age. Consistent with two large epidemiological studies of >40,000 individuals (Biino et al., 2013; J. Zhang, Li, & He, 2015), we found that platelet count decreased by ~6% for each decade of life (Figure 3b-c). This reflects a decline of 24% between the ages of 20 and 60, although the effect sizes vary by cohort and our estimate is likely overestimated due to the small size of our cohort. As expected, total platelet count tended to be consistently *positively* correlated with mtDNAcn, CS and RC activities in PBMCs (r=0.031-0.38) (Figure 3d). Therefore, the age-related loss of platelets (and of the mtDNA contained within them) could account for the previously reported age-related decline in mtDNAcn from studies using either whole blood (Mengel-From et al., 2014; Verhoeven et al., 2018) (which includes all platelets) or PBMCs (R. Zhang, Wang, Ye, Picard, & Gu, 2017) (which include fewer contaminating platelets).

We directly tested this hypothesis by immunodepleting platelets from “clean” PBMCs and comparing three resulting fractions: total PBMCs, actively platelet-depleted PBMCs, and platelet-enriched eluate. As expected, platelet depletion decreased mtDNAcn, CS, and RC activities, indicating that contaminating platelets exaggerated specific mitochondrial features by 9-22%, except for complex IV (Figure 3f-g). Moreover, the platelet-enriched eluate showed 23-100% higher mitochondrial activities relative to total PBMCs, providing direct evidence that the active platelet depletion method was effective and that platelets inflate estimates of mitochondrial abundance and RC activity in standard PBMCs prepared with two passive platelet-depletion steps. Interestingly, the composite MHI was minimally affected by the platelet depletion procedure, suggesting that this multivariate index of respiratory chain capacity on a per-mitochondrion basis may be more robust to platelet contamination than its individual features

### Individual cell subtypes are biologically distinct from PBMCs contaminated with platelets

Mitochondrial phenotyping was performed in FACS-purified immune cells, in parallel with PBMCs. To obtain sufficient numbers of cells for mitochondrial phenotyping, we selected the 6 most abundant cell subtypes for each individual and isolated 5×10^6^ cells for each sample. Because memory subtypes were relatively rare, central and effector memory (CM and EM) subtypes were pooled for CD4^+^ and/or CD8^+^ (CM-EM). This generated a total of 340 biological samples, including 136 biological replicates, yielding 204 individual participant/cell subtype combinations used in our analyses.

Among cell subtypes, CS activity was highest in monocytes and B cells, and lowest in CD4^+^ naïve T cells, with other cell types exhibiting intermediate levels (Figure 4a). Regarding mitochondrial genome content, B cells had the highest mtDNAcn with an average 451 copies per cell compared to neutrophils and NK cells, which contained only an average of 128 (g=5.94, p<0.0001) and 205 copies (g=3.84, p<0.0001) per cell, respectively (Figure 4b). Naïve and memory CD4^+^ and CD8^+^ T lymphocytes had intermediate mtDNAcn levels of ~300 copies per cell, except for CD8^+^ naïve cells (average of 427 copies per cell). Between cell types, CS activity and mtDNAcn differed by up to 3.52-fold.

**Figure 4.**
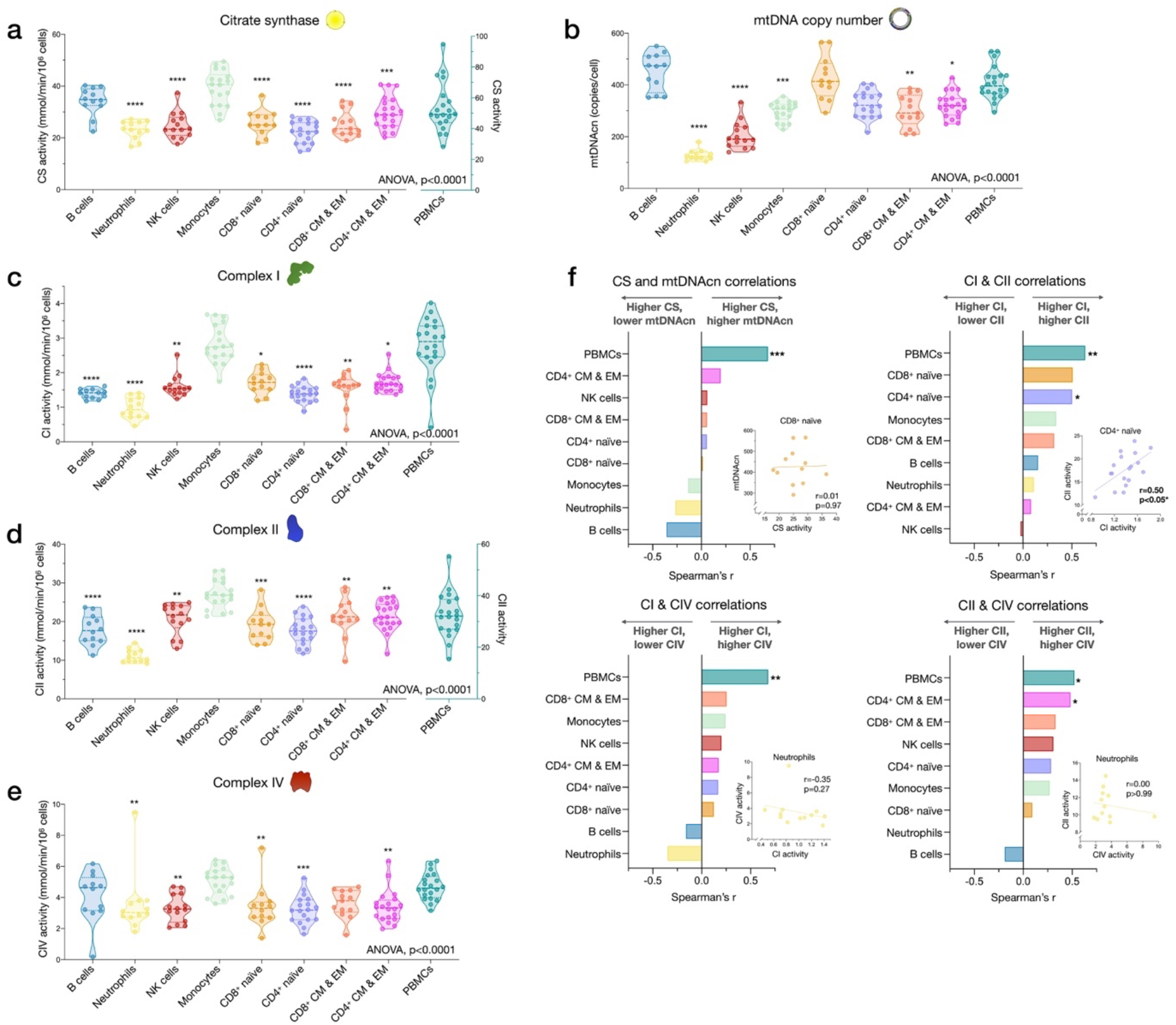
Cell subtype differences in mitochondrial content and RC function. (**a-e**) Violin plots illustrating immune cell type differences in mitochondrial features across cell subtypes and total PBMCs. For each individual, only the 6 most abundant cell types were analyzed (n=21 individuals, 12-18 per cell subtype). Dashed lines are median (thick) and 25th and 75th quartiles (thin). P-values from One-Way non-parametric ANOVA Kruskal-Wallis test, post-hoc Dunn’s multiple comparisons relative to PBMCs. (**f**) Spearman’s r inter-correlations of mitochondrial features across subtypes. Insets show the scatterplots for selected correlations. p<0.05*, p<0.01**, p<0.001***, p<0.0001****.

In relation to RC function, monocytes had the highest complex I, II, and IV activities. Consistent with their low mtDNAcn, neutrophils also had the lowest activities across complexes, whereas naïve and memory subtypes of T and B lymphocytes presented intermediate RC enzyme activities (Figure 4c-e). PBMCs had up to 2.9-fold higher levels of CS, CI, and CII activity per cell than any of the individual cell subtypes measured, again consistent with platelet contamination.

The correlations between different mitochondrial features indicate that CS and mtDNAcn were only weakly correlated with each other, and in some cases were negatively correlated Figure 4f). This finding may be explained by the fact that although both CS and mtDNAcn are positively related to mitochondrial content, CS may be superior in some cases (Larsen et al., 2012), inadequate in specific tissues (McLaughlin et al., 2020), and that mtDNAcn can change independently of mitochondrial content and biogenesis (Picard, 2021). For RC complexes CI, CII, and CIV, which physically interact and whose function is synergistic within the inner mitochondrial membrane, correlations tended to be positive, as expected (Figure 4f). However, relatively weak and absent inter-correlations between mitochondrial features in some cell types reveal that each metric (i.e., content features and enzymatic activities) provides relatively independent information about the immune cell mitochondrial phenotype.

We extended the analyses of univariate metrics of mitochondrial content and function by exploring the multivariate MHI, which significantly differed between cell subtypes (p<0.0001) (Figure 4-figure supplement 1). These results are discussed in Appendix 2.

### Mitochondrial features exhibit differential co-regulation across immune cell subtypes

Next, we asked to what extent mitochondrial markers correlate across cell subtypes in the same person (co-regulation). For example, we determined whether having high mtDNAcn or low MHI could constitute coherent properties of an individual that are expressed ubiquitously across cell types (e.g., an individual with the highest mtDNAcn in B cells also having the highest mtDNAcn in all cell types), or if these properties are specific to each cell subtype.

CS activity and mtDNAcn were moderately co-regulated across cell subtypes (average correlation r_z’_=0.63 and 0.53, respectively) (Figure 4-figure supplement 1c-d). In comparison, RC enzymes showed markedly lower correlations between cell types and some cell types were not correlated with other cell types, revealing a substantially lower degree of co-regulation among RC components than in mitochondrial content features. MHI showed moderate and consistent positive co-regulation across cell types (average r_z’_=0.37). Notably, PBMCs exhibited moderate to no correlation with other cell subtypes, further indicating their departure from purified subtypes (Figure 4-figure supplement 2). Together, these subtype-resolution results provide a strong rationale for performing cell-type specific studies when examining the influence of external exposures and person-level factors on immune cells’ mitochondrial bioenergetics, including the influence of sex and age.

### Mitochondrial content and RC function differ between women and men

To explore the added value of cell subtype specific studies when applied to real-world questions, we systematically compared CS activity, mtDNAcn, RC activity, and MHI between women and men (Figure 5a-f). Given the exploratory nature of these analyses with a small sample size, only two results reached statistical significance (analyses non-adjusted for multiple testing). Compared to men, women had 29% higher CS activity in CD8^+^ CM-EM T cells (g=1.52, p<0.05, Figure 5a) and 26% higher CI activity in monocytes (g=1.35, p<0.05, Figure 5c). Interestingly, across all cell subtypes examined, women showed a trend (p=0.0047, Chi-square) for higher CS activity (range: 4-29%, g=0.20-1.52, Figure 5a) and higher CII activity than men (range:1-10%, g=0.03-0.56, Figure 5d).

**Figure 5.**
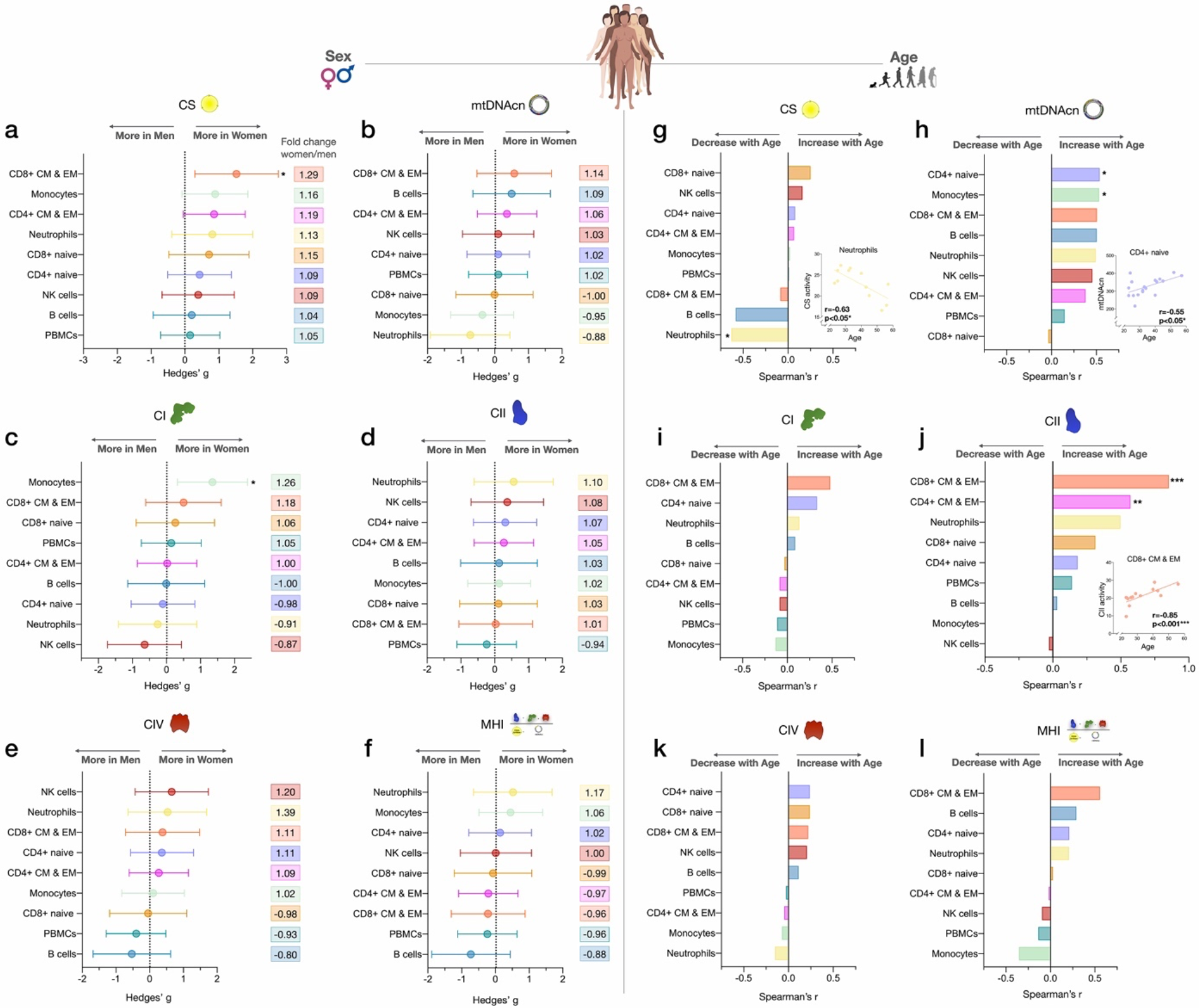
Associations of mitochondrial features with sex and age across cell subtypes. (**a-f**) Effect size of sex differences in mitochondrial activity across cell subtypes quantified by Hedges’ g. The fold change computed from raw values is shown on the right. P-values from Mann-Whitney test, not adjusted for multiple comparisons. Error bars reflect the 95% C.I. on the effect size. (**g-l**) Association of age and mitochondrial features across cell subtypes. P-values from Spearman’s r correlations, not adjusted for multiple comparisons. n=21 (11 women, 10 men), p<0.05*, p<0.01**, p<0.001***.

Other notable trends requiring validation in larger cohorts suggested that compared to women, men exhibited higher mtDNAcn in monocytes and neutrophils (range: 5-12%, g=0.37-0.73, Figure 5b), higher CI activity in neutrophils and NK cells (range: 9-13%, g=0.26-0.64, Figure 5c), and higher CIV activity specifically in B cells (20%, g=0.53, Figure 5e). Cells exhibiting the largest degree of sexual dimorphism on the integrated MHI were neutrophils (17% higher in women, g=0.52) and B cells (12% higher in men, g=0.73) (Figure 5f). In contrast, none of these differences were detectable in PBMCs, further illustrating the limitation of mixed cells to examine sex differences in mitochondrial function.

### Age associations with mitochondrial content and RC function

We then explored the association between mitochondrial features and age. With increasing age, CS activity was relatively unaffected except in neutrophils, where it significantly decreased by ~7% per decade (r=−0.63, p<0.05) (Figure 5g). In comparison, the correlation between age and mtDNAcn was positive among 7 out of 8 cell subtypes, with the exception being CD8^+^ naïve T cells. Interestingly, CD8^+^ naïve T cells is the cell type that exhibited the strongest age-related decline in abundance. CD4^+^ naïve T cells and monocytes showed the largest age-related change in mtDNAcn, marked by a significant ~10% increase in the number of mitochondrial genome copies per cell per decade of life (r=0.54, p<0.05 for both, Figure 5h).

For RC function, an equal number of cell subtypes with either positive or negative correlations with age were found, except for CII (Figure 5i-l). Due to our small sample size, only CD4^+^ and CD8^+^ CM-EM T cells CII activity significantly increased with age (r=0.57 and 0.85, p<0.01 and 0.001 respectively, Figure 5j). However, CII activity was positively correlated with age across all cell types except for monocytes and NK cells. In contrast, CI and CIV activities were only weakly associated with age, highlighting again differential regulation and partial “biological independence" of different RC components. Of all cell types, CD8^+^ CM-EM T cells showed the most consistent positive associations for all RC activities and age, most notably for CII where the enzymatic activity per cell increased a striking ~21% per decade (r=0.85, p<0.001).

Overall, these exploratory data suggest that age-related changes in CS activity, mtDNAcn, and RC function are largely cell-type specific. This conclusion is further reinforced by analyses of PBMCs where mitochondrial features consistently did not significantly correlate with age (r=0.008-0.15, absolute values) (Figure 5g-l). Larger studies adequately powered to examine sex- and age-related associations are required to confirm and extend these results.

### Cell subtype distributions exhibit natural week-to-week variation

Samples collected weekly over 9 weeks from one repeat participant were used to i) examine the stability of cell-type specific mitochondrial phenotypes described above, and ii) quantify whether and how much mitochondrial content/function change over time (Figure 6a). First focusing on immune cell distribution, the cell subtype with the least week-to-week variation in abundance was CD8^+^ EM (root mean square of successive differences [rMSSD]=0.22, coefficient of variation [C.V.]=19.5%), which varied between 6.3% (week 2, highest) and 3.3% of all circulating cells (week 9, lowest) (Figure 6-figure supplement 1). Other subtypes such as CD4^+^ TEMRA (min=0.02% to max=0.62%) and neutrophils (min=3.9% to max=31.8%) varied week-to-week by an order of magnitude (i.e., 10-fold), similar to the between-person variation among the cohort (see Figure 1e and Supplementary file 2). The circulating abundance of B cells varied by up to 1.1-fold (min=0.86% to max=1.8%). Together, these time-course results illustrate the dynamic remodeling of circulating leukocyte populations (and therefore PBMC composition) within a single person.

**Figure 6.**
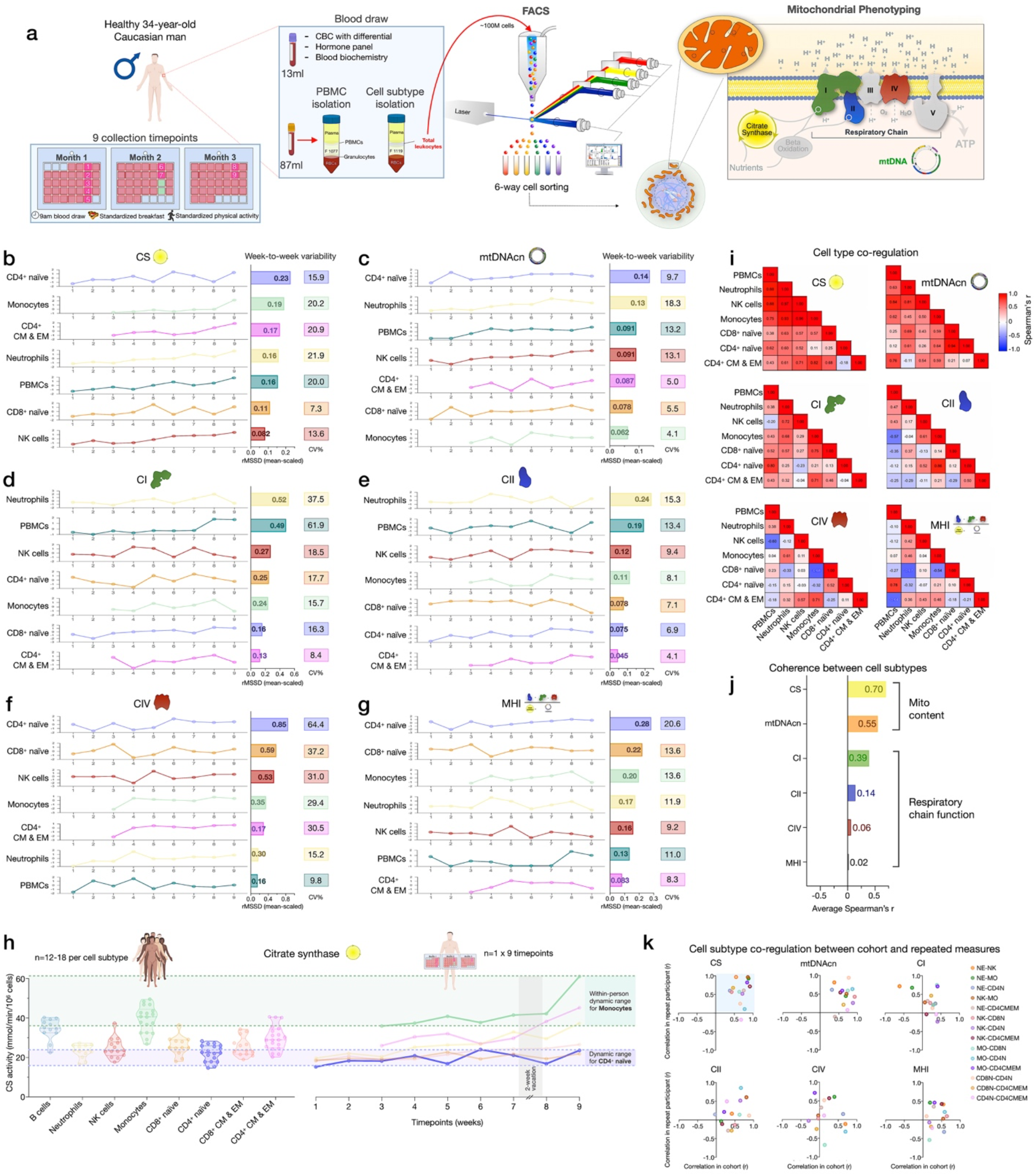
Within-person variability of mitochondrial features across cell subtypes. (**a**) Overview of the repeat participant design, including blood collection, processing, and analysis. All samples were collected, stored, and processed as a single batch with samples from the cohort. Mitochondrial phenotyping platform schematic adapted from Figure 1A (Picard et al., 2018). (**b-g**) Natural weekly variation for each mitochondrial feature across cell subtypes in the same person across 9 weeks represented as scaled centered data where 1 unit change represents a one-standard deviation (S.D.) difference. Root mean square of the successive differences (rMSSDs) quantify the magnitude of variability between successive weeks. The coefficients of variation (C.V.) quantify the magnitude of variability across all time points. Monocytes and CD4^+^ CM-EM were not collected on weeks 1 and 2. (**h**) Side-by-side comparison of CS activity between the cohort (n=12-18 per cell subtype) and the repeat participant (n=7-9 time points) across cell subtypes. The dynamic range of two cell subtypes are highlighted: monocytes and CD4^+^ naïve T cells. (**i**) Within-person correlation matrices between cell subtypes for each mitochondrial feature over 9 weeks, illustrating the magnitude of correlation (co-regulation) between cell subtypes. (**j**) Average inter-correlation across all cell subtypes by mitochondrial feature (calculated using Fisher z-transformation) indicating the degree of coherence within-person. (**k**) Comparison of co-regulation patterns among mitochondrial features between the cohort and the repeat participant. Each datapoint represents a cell subtype pair, indicating moderate agreement (datapoints in top right quadrant).

The correlations between immune cell type composition at each week and PBMC mitochondrial features are shown in Figure 6-figure supplement 2. On weeks when the participant had higher circulating levels of EM and TEMRA CD4^+^ and CD8^+^ lymphocytes, most mitochondrial features were considerably lower in PBMCs. The associations between CBC-derived cell proportions and PBMCs mitochondrial features tended to be weaker and in opposite direction at the within-person level compared to the cohort (Figure 6-figure supplement 2c-d), but again document the influence of cell type composition on PBMC mitochondrial phenotypes.

### Mitochondrial content, mtDNAcn and RC activity exhibit natural week-to-week variation

The 6 most abundant cell subtypes analyzed for this individual included: neutrophils, NK cells, monocytes, naïve and CM-EM subtypes of CD4^+^ T cells, and naïve CD8^+^ T cells. The robust cell type differences in mitochondrial content and RC activities reported above in our cohort were conserved in the repeat participant. This includes high mtDNAcn in CD8^+^ naïve T cells (average across 9 weeks=400 copies/cell, 427 in the cohort) and lowest mtDNAcn in neutrophils (average=123 copies/cell, 128 in the cohort).

All mitochondrial metrics exhibited substantial weekly variation across the 9 time points. Different cell types showed week-to-week variation in CS, mtDNAcn, and RC activity ranging from 4.1 to 64.4% (Figure 6b-f). In most cases, the observed variation was significantly greater than the established technical variation in our assays (see Supplementary file 3), providing confidence that these changes in mitochondrial content and function over time reflect real biological changes rather than technical variability.

We then asked how much the same metrics naturally vary within a person relative to differences observed between people in the heterogeneous cohort. Remarkably, the 9-week range of natural variation within the same person was similar to the between-person differences among the cohort. Figure 6h and Figure 6-figure supplement 3 provide a side-by-side comparison of the cohort and repeat participant mitochondrial features (CS, mtDNAcn, RC activities, and MHI) on the same scale. A similar degree of variation (9.8-61.9%) was observed in PBMCs (Figure 6b-g), although again this variation may be driven in large part by variation in cell composition.

And as in the cohort, CS and mtDNAcn were also the most correlated across cell types (average r_z’_=0.55-0.70) (Figure 6i-k, Figure 6-figure supplement 2e), indicating partial co-regulation of mitochondrial content features across cell subtypes.

### Mitotypes differ between immune cell subtypes

To examine cell type differences more fully, and in line with the concept of mitochondrial functional specialization, we performed exploratory analyses of cell subtype-specific mitochondrial phenotypes, or *mitotypes*, by mathematically combining multiple mitochondrial features in simple graphical representations (listed and defined in Figure 7-figure supplement 1). Each mitotype can be visualized as a scatterplot with two variables of interest as the x and y axes. Cell types that align diagonal from the origin of the plot indicate the same mitotype profile (Figure 7a).

**Figure 7.**
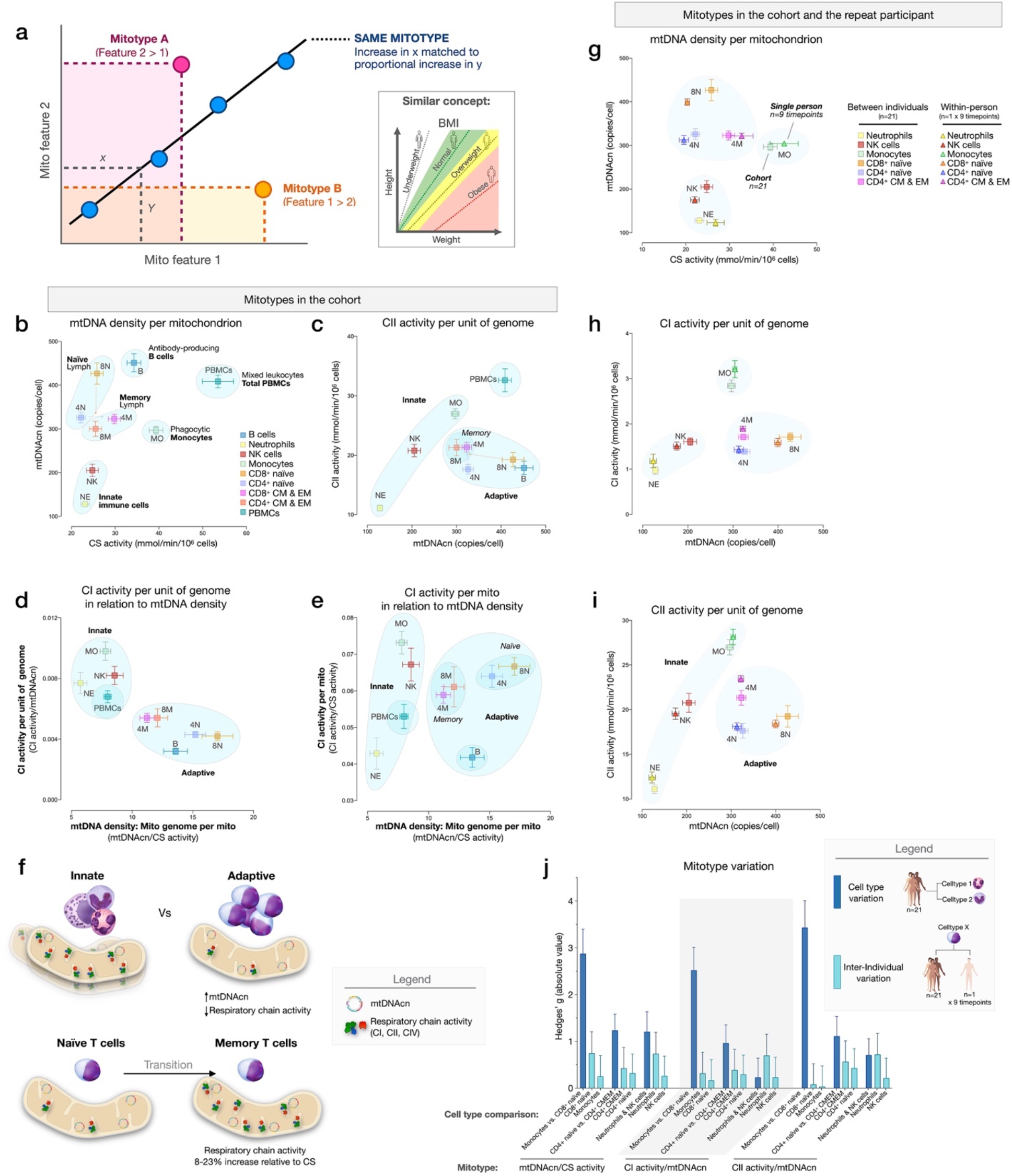
Mitotypes in purified leukocyte populations from the cohort and repeated-measures. (**a**) Schematic illustrating the multivariate approach to generate and visualize mitotypes by putting into relation two or more mitochondrial features. Note the similarity and added insight relative to single metrics, similar to the integration of height and weight into the body mass index (BMI). (**b-e**) Selected mitotypes plotted for each cell subtype among the cohort. Data are means ± SEM (n=12-18). Overlaid shaded areas denote general leukocyte categories, for visualization purposes only. (**f**) Summary of mitotype differences between (i) innate vs adaptive subdivisions, and (ii) naïve vs memory T cells. (**g-i**) Validation of subtype-specific mitotype differences in the repeat participant, illustrating the conserved nature of mitotypes across individuals. Only the six cell subtypes analyzed in the repeat participant are plotted. Data are means ± SEM (n=7-9 for the repeat participant, 12-18 for the cohort). (**j**) Comparison of the magnitude of the difference (Hedges’ g) in mitotypes between cell types, and between individuals. Dark blue bars indicate the magnitude of the dominant difference in mitotypes between cell subtypes. Light blue bars indicate the magnitude of the difference between the cohort and the repeat participant within a cell type. Error bars are the 95% C.I. of the effect size.

The first mitotype examined mtDNA copies per cell (mtDNAcn) relative to mitochondrial content per cell (CS activity), creating a mitotype (mtDNAcn/CS) reflecting *mtDNA density per mitochondrion* (Figure 7b). Alone, the mtDNA density per mitochondrion mitotype provided remarkable separation of cell subtypes. Neutrophils and NK cells were low in both mtDNAcn and CS activity, B cells were high on both metrics, monocytes had the lowest mtDNA density, whereas CD8^+^ naïve T cells exhibited the highest mtDNA density of all cell subtypes tested. Figure 7c-e illustrate other mitotypes including i) CII activity per unit of mitochondrial genome (CII/mtDNAcn), as well as more complex combinations of variables such as ii) CI activity per mtDNA (CI/mtDNAcn ratio on y axis) in relation to mtDNA density (mtDNAcn/CS activity on x axis), and iii) CI activity per mitochondrial content (CI/CS, y axis) in relation to mtDNA density (mtDNAcn/CS, x axis). As Figure 7b-e shows, PBMCs generally exhibit a similar mitotype as innate immune cell subtypes (monocytes, NK cells, and neutrophils on the same diagonal), and are relatively distinct from lymphocyte subpopulations.

This mitotype-based analysis revealed two main points. First, cells of the innate and adaptive immune subdivisions contain mitochondria that differ not only quantitatively in their individual metrics of mitochondrial content and RC activity, but also qualitatively, as illustrated by the distinct clustering of neutrophils, monocytes, and NK cells (innate) within similar mitotype spaces, and the distinct clustering of all lymphocyte subtypes together in a different space. Compared to cells of the innate immune compartment, lymphocytes (adaptive) had higher mtDNAcn and lower respiratory chain activity. Second, compared to naïve subsets of CD4^+^ and CD8^+^ T cells, which themselves have relatively distinct mitotypes (e.g., CII/mtDNAcn, Figure 7c), both memory CD4^+^ and CD8^+^ subtypes converged to similar mitotype spaces. Functionally, this naïve-to-memory transition is well known to involve a metabolic shift including changes in spare respiratory capacity, mitochondrial content, and glucose and fatty acid uptake (Nicoli et al., 2018; van der Windt et al., 2012). The mitotype analysis showed that compared to naïve cell subtypes, memory subtypes exhibit 26-29% lower mtDNA density per mitochondrion, but an 8-23% increase in RC activity per mitochondrion in CD4^+^ T cells, although not in CD8^+^ T cells (Figure 7f).

### Stability of mitotypes

The 6 cell subtypes analyzed in the repeat participant and the matching cell types for the cohort showed a high degree of agreement when plotted on the same mitotype plots. Again, cell types belonging to the innate and adaptive immune subdivisions clustered together, and naïve and memory subtype differences were similarly validated at the within-person level (Figure 7g-i), demonstrating the conserved nature of immune cell mitotypes in our sample. On average, the magnitude of variation between cell subtypes (e.g., monocytes vs neutrophils) was 12.5-fold larger than the differences between the cohort and the repeat participant, indicating that immune cell subtypes have conserved mitotypes, exhibiting relative stability across individuals.

### Evidence for a sex- and age-related bias in mitotypes

We next sought to systematically examine if mitotypes differ between women and men. Mitotypes were organized into five categories of indices based upon their features, yielding a total of 16 mathematically-distinct mitotypes (see Figure 7-figure supplement 1). For each mitotype, we quantified the magnitude of the difference between women and men by the effect size (g), ranked all mitotype x cell subtype combinations (16 mitotypes x 9 cell subtypes), and analyzed the distribution of these indices by sex. The majority of mitotypes reflecting mitochondrial RC activities per CS activity were higher in men (p<0.0001, Chi-square), while RC activity per mtDNA density (p<0.001) and RC activity per genome in relation to mtDNA density mitotypes (p<0.01) were predominantly higher in women (Figure 8a). The magnitude of sex differences ranged from 17% higher in men (CI/CS in CD4^+^ CM-EM T cells, g=1.14) to 38% higher in women (CII/mtDNA density in neutrophils, g=1.37) (Figure 8b). The direction of sex differences for all mitotypes (e.g., higher in women or in men) with effect sizes is illustrated in Figure 8c. The average effect size across all mitotypes was 0.31 (small) in CD4^+^ naïve T cells, compared to monocytes where the average effect size was 0.71 (medium). Compared to purified cell subtypes, the magnitude of sex differences in PBMCs was blunted.

**Figure 8.**
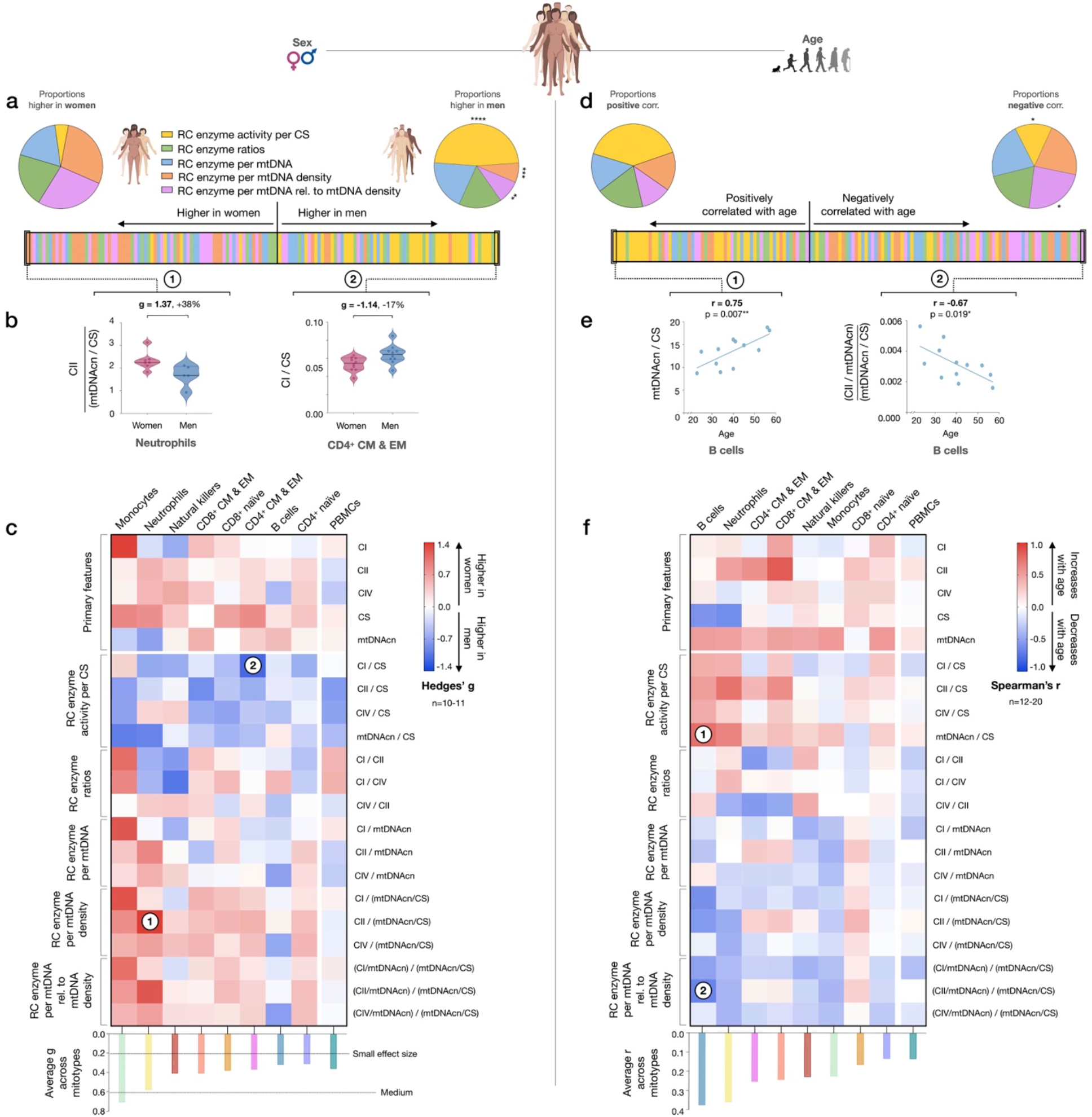
Mitotype distribution and strength of difference across sex and age. (**a**) Ranking of mitotype indices by the effect size (Hedges’ g) between women and men. A total of 16 mitotype indices were computed, subdivided into 5 main color-coded categories (see Figure 7-figure supplement 1). Pie charts illustrate the proportion mitotypes belonging to each category that are either higher in women (*left*) or in men (*right*). P-values for enrichment of sexually dimorphic mitotypes are derived from Chi-square test. (**b**) Violin plots illustrating the two mitotypes with the largest sex differences, both showing large effect sizes (g). (**c**) Heatmap of sex differences for primary measures of mitochondrial function (*top*) and multivariate mitotypes (*bottom*) across cell subtypes. The histogram at the bottom shows the average effect size across all mitotypes (calculated from absolute g values). (**d**) Ranking of mitotype indices by the strength and direction of their association with age, with enrichment analysis analyzed as for sex (Chi-square test). (**e**) Spearman’s r correlations of mitotypes/cell type combinations with the strongest positive and negative associations with age. (**f**) Heatmap of the age correlations (Spearman’s r) for primary features and composite mitotypes across cell subtypes. The histogram (*bottom*) shows the average effect size (r) for each cell subtype (calculated using absolute values and Fisher z-transformation). p<0.05*, p<0.01**, p<0.001***, p<0.0001****.

Using the same approach, we then systematically quantified the relationship between mitotypes and age. Mitotypes reflecting RC activity per CS activity were predominantly positively correlated with age (p<0.05), while RC activities per genome in relation to mtDNA density were generally negatively correlated with age (p<0.05) (Figure 8d). This finding is consistent with the overall age-related increase in mtDNAcn across cell subtypes, and could indicate a general decrease in the RC output per unit of mitochondrial genome with aging in immune cells. The strength of these correlations ranged from r=−0.67 to 0.75 (Figure 8e). The correlations of individual mitotypes with age for each cell subtype are shown in Figure 8f. Again, PBMCs showed among the weakest associations with either sex or age (Figure 8c and f). Thus, even if specific cell subtypes reveal consistent sex- and age-related differences, PBMCs offer modest to no sensitivity to detect these associations.

### Associations of blood biomarkers with subtype-specific mitochondrial features

Finally, to explore the source of inter-individual differences and within-person dynamics over time described above, we asked to what extent subtype-specific mitochondrial features were correlated with standard blood biomarkers, including a panel of sex hormones, inflammatory markers, metabolic markers, and standard clinical blood biochemistry (Figure 9a).

**Figure 9.**
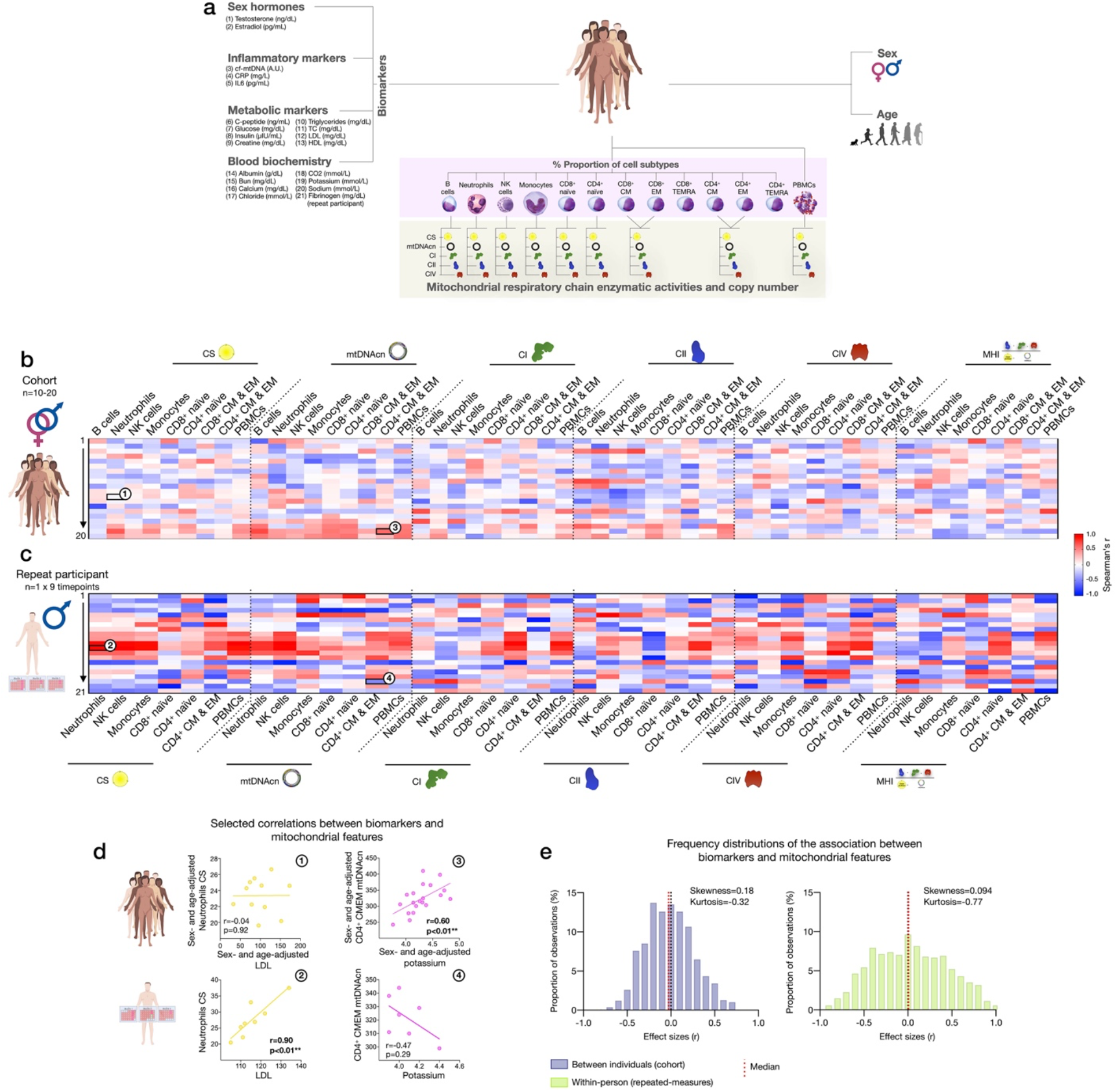
Association of blood biomarkers with mitochondrial parameters across cell subtypes and primary mitochondrial features. (**a**) Overview of blood biochemistry, hormonal, and metabolic biomarkers collected for each participant. (**b**) Sex- and age-adjusted correlations between blood biomarkers and mitochondrial features across cell subtypes for the cohort (n=10-20 per mito:biomarker combinations) shown as a heatmap. (**c**) Same as (b) using weekly measures of both mitochondrial features and biomarkers in the repeat participant. (**d**) Scatterplots of the indicated correlations between Neutrophils CS activity and LDL cholesterol (*left*), and CD4^+^ CM-EM mtDNAcn and potassium (K^+^) (*right*) for the cohort (*top row*) and the repeat participant (*bottom row*). (**e**) Frequency distributions of the aggregated effect sizes between biomarkers and mitochondrial features across cell subtypes for the cohort (total correlation pairs=1,080) and the repeat participant (total correlation pairs=882).

At the cohort level, sex- and age-adjusted partial correlations between blood biomarkers and cell subtype mitochondrial phenotypes were relatively weak (average absolute values rz’=0.23, Figure 9b), indicating that circulating neuroendocrine, metabolic and inflammatory factors are unlikely to explain a large fraction of the variance in inter-individual differences in mitochondrial biology. At the within-person level, week-to-week variation is independent of constitutional and genetic influences; additionally, behavior (e.g., levels of physical activity, sleep patterns, etc.) is more stable relative to between-person differences among a cohort of heterogenous individuals. Accordingly, compared to the cohort, the strength of biomarker-mitochondria associations was on average r_z’_=0.39, or 70% larger in the repeat participant than the cohort (p<0.0001, comparing absolute correlation values) (Figure 9c). In particular, lipid levels including triglycerides, total cholesterol, and low- and high-density lipoproteins (LDL, HDL) were consistently positively correlated with markers of mitochondrial content (CS activity and mtDNAcn), with the largest effect sizes observed among innate immune cells: neutrophils, NK cells, and monocytes (Figure 9c, red area on the heatmap). In these cells, lipid levels accounted on average for 53% of the variance (r^2^) in CS activity and 47% in mtDNAcn, possibly reflecting an effect of lipid signaling on mitochondrial biogenesis (Iershov et al., 2019; Lindquist et al., 2018; Picard et al., 2012; Turner et al., 2007). We also note major divergences in the correlation patterns between the cohort and repeat participant (Figure 9d-e). Although limited by the small number of observations, these results highlight the value of repeated-measures study designs to examine the influence of metabolic and other humoral factors on human immune mitochondrial biology which may be blunted at the group-level (Fisher, Medaglia, & Jeronimus, 2018).

## Discussion

Developing approaches to quantify bioenergetic differences among various tissues and cell types is critical to define the role of mitochondria in human health and disease. Here we isolated and phenotyped multiple molecularly-defined immune cell subtypes and mixed PBMCs in >200 person-cell type combinations among a diverse cohort of women and men, and in repeated weekly measures in the same participant. Our biochemical and molecular results confirm and extend previous knowledge of bioenergetic differences across human immune cell types established using extracellular flux analysis, protein and mtDNA quantification (Chacko et al., 2013; Lee et al., 2021; Maianski et al., 2004; Pyle et al., 2010). We also report preliminary evidence that mitochondrial phenotypes vary with age and sex, which PBMCs lack the sensitivity to detect. Importantly, our results confirm and quantify the extent to which PBMCs mitochondrial measurements are confounded by i) cell type composition, ii) platelet contamination, iii) mitochondrial properties across different cell subtypes, and iv) by the dynamic remodeling of cell type composition and bioenergetics over time. In addition, large week-to-week, within-person variation in both cell subtype proportions and mitochondrial behavior points to heretofore underappreciated dynamic regulation of mitochondrial content and function over time in humans. Overall, our results provide standardized effect sizes of mitochondrial variation in relation to multiple key covariates, highlights the value of repeated-measures designs to carefully examine the mechanisms regulating mitochondrial health in humans, and call for the replication and extension of these findings in larger cohorts.

This study emphasizes the value of using purified cell populations over PBMCs for mitochondrial analyses. In many cases, associations with moderate to large effect sizes in specific cell subtypes were either not observed or blunted in PBMCs. For example, we found no correlation between age and PBMC MHI neither in this study nor in a previous one (Karan et al., 2020), whereas purified cell subtypes showed associations. Interestingly, in the mitotype plots, total PBMCs had similar mitotypes (along the same mitotype diagonal space) as cells of the innate subdivision, namely neutrophils, NK cells, and monocytes. If PBMCs were composed uniquely of a mixture of lymphocytes and monocytes, the natural expectation is that PBMCs would lie somewhere between the specific subsets that compose it. Instead, PBMCs occupy an entirely different and unexpected mitotype space. Additionally, our platelet depletion experiment leaves little doubt that platelet contamination skews the measurements of several mitochondrial features in PBMCs, with some features being apparently more affected than others, and yielding contradictory results: for example, PBMCs have higher CS activity values than any of the constituent cells (see Figure 4a). Although we cannot entirely rule out potential contamination of individual cell types with residual platelets, the FACS labeling, washing, and sorting procedures must produce the purest sample with the highest degree of biological specificity.

A major frontier for the human immunometabolism field consists in defining temporal trajectories of change in specific cell types (Artyomov & Van den Bossche, 2020). Achieving this goal promises to transform knowledge of immune and mitochondrial biology and allow for rational design of therapeutic approaches for immunometabolic conditions (Artyomov & Van den Bossche, 2020). A primary finding from our analyses is the natural within-person variation in mitochondrial features, providing initial insight into the temporal dynamics of immunometabolism in human leukocytes. Sorting immunologically-defined cell subtypes removed the potential confound of week-to-week changes in cell type distributions, an inherent confounding variable in PBMCs, and therefore adds robustness to our observations. Mitochondrial features within immune cells exhibited state-like properties that varied by >20-30% week-to-week, warranting future, adequately-powered studies of causal influences. Previously, in PBMCs, up to 12% of the inter-individual variation in MHI was found to be attributable to positive mood (e.g., excited, hopeful, inspired, love) the night prior to blood draw (Picard et al., 2018), implying that psychosocial factors could in part contribute to dynamic variation in leukocyte mitochondrial function over 12-48 hours. However, limitations in this prior study – including the use of PBMCs and a single measurement time point for MHI – call for additional studies to disentangle the independent contributions of behavioral, psychosocial, nutritional, and other factors on specific mitochondrial features. Importantly, mitochondrial changes in the present study took place within less than one week. Tracking dynamic changes in biological markers with high intra-individual variability, such as cortisol (Segerstrom, Sephton, & Westgate, 2017), necessarily requires repeated-measures designs with sufficient temporal resolution. Therefore, the present results and others show that establishing the exact temporal dynamics of leukocyte mitochondrial variations, and immunometabolism in general, will require repeated assessments with substantially greater temporal resolution than weekly measures.

Animal studies have consistently identified sexually dimorphic mitochondrial features, such as greater mitochondrial content in females (reviewed in (Ventura-Clapier, Piquereau, Veksler, & Garnier, 2019)). Likewise, in humans, PBMCs from women had greater CS activity and greater CI and CII-mediated respiration (Silaidos et al., 2018). Our data show similar changes in enzymatic activities for most, but not all, cell types, suggesting that the magnitudes of sex differences are likely cell-type specific. Therefore, methods offering a sufficient level of biological specificity deployed in adequately powered samples are needed to reproducibly and accurately quantify sex differences among immune cell mitochondria. We also note that the binary definition of sex used in this study (i.e., sex assigned at birth) paints a rather incomplete picture. Exploring the interplay between mitochondria, sex and gender in humans will require more refined approaches that consider a range of biological characteristics (e.g., hormones, chromosomes, anatomy), alongside gender identity and its proceeding gendered exposures (e.g., differential diet or involvement in physical activity depending on gender/sex) (Fausto-Sterling, 2005; Johnson & Repta, 2012; Ritz et al., 2014).

In relation to age, age-related decline in mtDNAcn has consistently been reported in whole blood (Mengel-From et al., 2014; Verhoeven et al., 2018), PBMCs (R. Zhang et al., 2017), and skeletal muscle tissue (Hebert et al., 2015; Short et al., 2005), although not in liver (Wachsmuth, Hübner, Li, Madea, & Stoneking, 2016). However, the interpretation of these results must take into account the existence of cell mixtures and platelet contamination, particularly for blood and PMBCs (Banas et al., 2004; Urata et al., 2008). In one large adult cohort study, accounting for cell type distribution and platelet count through measurement and statistical adjustments eliminated initial associations between mtDNAcn and age (Moore et al., 2018), suggesting that the *apparent* age-related decline in mtDNAcn in human blood in fact reflect a change in blood composition (fewer platelets in older people, explaining why mtDNAcn appears lower). In purified immune cell subtypes from our small cohort, the opposite association was observed. Although based on whole-blood and tissue studies this finding was unexpected, it could be explained by well-known processes related to the quality of mtDNA (Picard, 2021). Mutations and deletions in mtDNA accumulate with age (Ye, Lu, Ma, Keinan, & Gu, 2014; R. Zhang et al., 2017), and mtDNA defects can trigger the compensatory upregulation of mtDNAcn to counteract that loss of intact mitochondria (Giordano et al., 2014; Yu-Wai-Man et al., 2010). Therefore, the observed positive correlation of cell type-specific mtDNAcn with age in our sample could reflect compensatory upregulation of mtDNA replication. Alternatively, this correlation could reflect impaired autophagic removal of mitochondria in aging cells, consistent with recent results suggesting that CD4^+^ T cells have impaired clearance of dysfunctional mitochondria (Bektas et al., 2019). Interestingly, the only cell type examined that did not exhibit positive correlation between mtDNAcn and age was CD8^+^ naïve T cells, which is also the only cell type whose abundance in circulation significantly declines with advancing age. The basis for the direction of this association requires further investigation.

Some limitations of this study must be noted. Although this represents, to our knowledge, the largest available study of mitochondrial biochemistry and qPCR in hundreds of human samples, the sample size of the cohort was small and the power to examine between-person associations was limited. Women and men were equally represented, but the sample size precluded stratification of all analyses by sex. Sex and age analyses are exploratory and findings need to be validated by future adequately powered studies. Likewise, the exhaustive repeated-measures design was carried out in only one participant and should be regarded as proof-of-concept. Additionally, because our mitochondrial phenotyping platform required ~5×10^6^ cells per sample, we could only collect the six most abundant cell subtypes from each participant, which in some instances reduced the final sample size for different cell subtypes. In order to accommodate the minimum cell number per sample, central and effector memory subtypes were pooled (CM-EM), although they may exhibit differences that are not resolved here. Furthermore, we recognize that additional cell surface markers may be useful to identify other cell populations (e.g., activated or regulatory lymphocyte subtypes). Finally, we did not test participants for cytomegalovirus (CMV) status, which could influence the proportion of immune cell subtypes.

Furthermore, our analysis focused on RC activity, which performs electron transport and generates the mitochondrial membrane potential (ΔΨm) across the inner mitochondrial membrane (Nicholls & Ferguson, 2013). Besides being used for ATP synthesis by complex V, RC activity and membrane potential also contributes to reactive oxygen species (ROS) production, calcium handling, and gene expression regulation, among other cellular processes (M. D. Brand, Chen, & Lehninger, 1976; Hernansanz-Agustín & Enríquez, 2021; Martínez-Reyes et al., 2016; Picard et al., 2014). Thus, the observed cell type differences in mitochondrial content or RC activities across human immune cell subtypes likely reflect not only cellular ATP demand, but also the unique immunometabolic, catabolic/anabolic, and signaling requirements among different immune cell subtypes that contribute to produce cell-specific mitotypes.

Overall, our study of mitochondrial profiling in circulating human immune cells filled three main knowledge gaps. First, it quantified confounds for PBMCs and showed how PBMCs fail to capture age- and sex-related mitochondrial recalibrations in specific immune cell populations, which is important for the design of future studies. Second, mitochondrial profiling precisely documented large-scale, quantitative differences in CS activity, mtDNAcn, and RC enzyme activities between well-known immune cell subtypes, contributing to our knowledge of the distinct metabolic characteristics among circulating immune cell types in humans. The qualitative and quantitative divergences were particularly emphasized by mitotypes, which highlighted conserved multivariate phenotypic features between lymphoid- and myeloid-derived immune cells, and between naïve and memory lymphocyte states. Third, this study documents potentially large week-to-week variation of mitochondrial activities that should be further examined in future studies. Together, this work provides foundational knowledge and a resource to develop interpretable blood-based assays of mitochondrial health.

## Methods

### Participants and Procedures

A detailed account of all methods and procedures is available in Appendix1. The study was approved by New York State Psychiatric Institute (Protocol #7618) and all participants provided written informed consent for the study procedures and reporting of results. Healthy adults between 20 and 60 years were eligible for inclusion. Exclusion criteria included severe cognitive deficit, symptoms of flu or other seasonal infection four weeks preceding the visit, involvement in other clinical trials, malignancy or other clinical condition, and diagnosis of mitochondrial disease. The main study cohort included 21 individuals (11 women, 10 men), with a mean age of 36 years (SD= 11, range: 23-57); there were 2 African Americans, 7 Asians, and 12 Caucasians. Morning blood samples (100 ml) were drawn between 9-10am from the antecubital vein and included one EDTA tube for complete blood count (CBC), two SST coated tubes for hormonal measures and blood biochemistry, and 11 Acid Dextrose A (ACD-A) tubes for leukocyte isolation and mitochondrial analyses. See Figure 1a-b for an overview of participants and procedures.

Additionally, repeated weekly measures were collected across 9 weeks from one healthy Caucasian man (author M.P., 34 years old) to assess within-person variability in mitochondrial measures and immune cell type distribution. To standardize and minimize the influence of nutritional and behavioral factors in the repeat participant, repeated measures were collected at the same time (9:00am), on the same day of the week (Fridays), after a standardized breakfast (chocolate crepe), ~30-60 minutes after a regular bicycle commute to study site.

### PBMCs and leukocyte isolation

A detailed version of the materials and methods is available in Appendix 1 of this article. Briefly, PBMCs were isolated on low density Ficoll 1077, and total leukocytes were separated on Ficoll 1119, centrifuged at 700 x g for 30 minutes at room temperature. Leukocytes were collected, washed, and centrifuged twice at 700 x g for 10 minutes to reduce platelet contamination. Pellets of 5×10^6^ PBMCs were aliquoted and frozen at −80°C for mitochondrial assays.

### Immunolabeling and fluorescence-activated cell sorting (FACS)

Antibody cocktails for cell counting (Cocktail 1) and cell sorting (Cocktail 2) were prepared for fluorescence-activated cell sorting (FACS). The following cell subtypes were identified: neutrophils, B cells, monocytes, NK cells, naïve CD4^+^ and CD8^+^, central memory (CM) CD4^+^ and CD8^+^, effector memory (EM) CD4^+^ and CD8^+^, and terminally differentiated effector memory cells re-expressing CD45RA (TEMRA) CD4^+^ and CD8^+^ (see Figure 1-figure supplement 1 and Supplementary file 1 for overview, and Supplementary file 4 for cell surface markers and fluorophores). A 2×10^6^ cell aliquot was labeled with Cocktail 1. The remainder of total leukocytes (~100×10^6^ cells) were incubated with Cocktail 2, washed, and used for FACS at final concentration of 20×10^6^ cells/ml.

Leukocytes were sorted using a BD™ Influx cell sorter using a 100 μm size nozzle. Sorting speed was kept around 11,000-12,000 events/second. Cell concentration for sorting was measured at about 15×10^6^ cells per ml. For each participant, 1×10^6^ cells (Cocktail 1 panel) were run first to calculate the potential yield of each subpopulation (total cell number x percentage of each subpopulation). The variable proportions of cell subtypes from person-to-person (provided in full in Supplementary file 2) determined which cell subtypes were collected from each participant, and 5×10^6^ cell aliquots of the six most abundant subpopulations were sorted. Purity checks were performed on all sorted subpopulations to ensure the instrument performance was good enough to reach the sorted population purity >95%. Data were processed using FCS Express 7 software (see Figure 1-figure supplement 2 for gating strategy).

### Mitochondrial enzymatic activities

Sorted cell subtypes were centrifuged at 2,000 x g for 2 minutes at 4°C and stored in liquid nitrogen (−170°C) for 4-12 months until mitochondrial biochemistry and mtDNAcn analyses were performed as a single batch. Cell pellets (5×10^6^) were mechanically homogenized with a tungsten bead in 500 ul of homogenization buffer as previously described (Picard et al., 2018) (see Appendix 1 for details).

Mitochondrial enzyme activities were quantified spectrophotometrically for citrate synthase (CS), cytochrome c oxidase (COX, Complex IV), succinate dehydrogenase (SDH, Complex II), and NADH dehydrogenase (Complex I) as described previously(Picard et al., 2018) with minor modifications described in Appendix 1. Each sample was measured in triplicates. Mitochondrial enzymatic activities were measured on a total of 340 samples (including 136 biological replicates), for a total of 204 unique participant-cell combinations. The technical variation for each enzyme, for each cell type, is detailed in Supplementary file 3.

### Mitochondrial DNA copy number

mtDNAcn was determined as described previously (Picard et al., 2018) using two different Taqman multiplex assays for ND1 (mtDNA) and B2M (nDNA), and for COX1 (mtDNA) and RnaseP (nDNA). mtDNAcn was calculated from the ΔCt obtained by subtracting the mtDNA Ct from nDNA Ct for each pair ND1/B2M and COX1/RNaseP, and mtDNAcn from both assays was averaged to obtain the final mtDNAcn value for each sample. The coefficients of variation (C.V.) for mtDNA for each cell subtype is detailed in Supplementary file 3 (average 5.1%).

### Platelet depletion in PBMCs

A further 9 community-dwelling older adults (mean age = 79, range: 64-89, 4 women and 5 men, including 7 white and 2 African American participants) were recruited for active platelet depletion experiments. Exclusion criteria included diseases or disorders affecting the immune system including autoimmune diseases, cancers, immunosuppressive disorders, or chronic, severe infections; chemotherapy or radiation treatment in the 5 years prior to enrollment; unwillingness to undergo venipuncture; immunomodulatory medications including opioids and steroids; or more than two of the following classes of medications: psychotropics, anti-hypertensives, hormones replacement, or thyroid supplements. Participants were recruited from a volunteer subject pool maintained by the University of Kentucky Sanders-Brown Center on Aging. The study was conducted with the approval of the University of Kentucky Institutional Review Board. Morning blood samples (20 mL) were collected by venipuncture into heparinized tubes. PBMCs were isolated from diluted blood by density gradient centrifugation (800 x g for 20 minutes) using Histopaque. Buffy coats were washed once, and cells were counted using a hemocytometer. PBMCs (20-30 M) were cryopreserved in liquid nitrogen in RPMI-1640 with 10% FBS and 10% DMSO until further processing.

For active platelet depletion, PBMCs were first thawed at room temperature and centrifuged at 500 x g for 10 minutes, counted and divided into 2 aliquots, including 10×10^6^ cells for total PBMCs, and 11×10^6^ cells for platelet depletion. The total PBMCs were centrifuged at 2,000 x g for 2 min at 4°C and frozen as a dry pellet at −80°C until processing for enzymatic assays and qPCR. The PBMCs destined for platelet depletion were processed immediately and depleted of platelets following manufacturer procedures using magnetically-coupled antibodies against the platelet marker CD61. This experiment yielded three samples per participant, including total PBMCs, platelet depleted PBMCs, and enriched platelets. Each sample was processed in parallel for RC enzymatic activity assays and mtDNAcn as described above.

### Statistical analyses

To adjust for potential order and batch effects across the 340 samples (31 samples per 96-well plate, 17 plates total), a linear adjustment was applied to raw enzymatic activity measures to ensure consistency between the first and last samples assayed. Samples from both the cohort and repeat participant were processed and analyzed as a single batch, ensuring directly comparable data.

Throughout, standardized effect sizes between cell subtypes and between groups were computed as Hedge’s g (g). Mann-Whitney T tests were used to compare sex differences in cell type proportions and mitochondrial measures. Spearman’s r (r) was used to assess strength of associations between continuous variables such as age and circulating proportions of cell subtypes. To assess to what extent mitochondrial features are correlated across cell subtypes (co-regulation) and to calculate the average correlation across mitotypes, Spearman’s r matrixes were first computed and transformed to Fisher’s Z’, and then averaged before transforming back to Spearman’s r (r_z’_). One-way non-parametric ANOVA Kruskal-Wallis with post-hoc Dunn’s multiple comparison tests were used to compare cell type mitochondrial measures in different cell subtypes and PBMCs. Between- and within-person variation were characterized using coefficients of variation (C.V.). The root mean square of successive differences (rMSSD) was computed to quantify the magnitude of variability between successive weeks for repeated measures. Chi-square tests were computed to compare proportion of mitotype indices categories (enzyme activity per CS, enzyme ratios, enzyme per mtDNA, enzyme per mtDNA density, and enzyme per mtDNA relative to mtDNA density) by age (lower vs higher with increased age) and sex (lower vs higher in men). Finally, one-way non-parametric Friedman tests with post hoc Dunn’s multiple comparisons were used to compare mitochondrial measures in platelet-depleted PBMCs, enriched platelets PBMCs, and total PBMCs. Statistical analyses were performed with Prism 8 (GraphPad, CA), R version 4.0.2 and RStudio version 1.3.1056. Statistical significance was set at p<0.05.

## Data Availability

All data generated and analyzed during this study, including mitochondrial biochemistry, mtDNA content, and blood chemistry, cell counts from CBC and flow cytometry, and de-identified participant information are included in the supporting data files. Source data files have been provided for Figures 1-9, and for figure supplements (Figure 1-figure supplement 3, Figure 2-figure supplement 1, Figure 4-figure supplement 1, Figure 4-figure supplement 2, Figure 6-figure supplement 1, Figure 6-figure supplement 2, Figure 6-figure supplement 3, and Supplementary file 3). Requests for resources or other information should be directed to and will be fulfilled by the corresponding author.

## Acknowledgements

Work of the authors is supported by the Wharton Fund and NIH grants MH119336, GM119793, MH122706, AG066828, AG056635, AG026307, and UL1TR001873. These studies used the resources of the Irving Cancer Center Core Facility funded in part through center grant P30CA013696.

## Supplemental Figure Legends

**Figure 1-figure supplement 1 – Diagram of leukocyte cell lineages.**

Figure 1-figure supplement 1 adapted from [Figure 3.34, OpenStax 2021, https://openstax.org/books/anatomy-and-physiology/pages/3-6-cellular-differentiation]. Cells analyzed in this study are circled in red. The cell surface markers used to define sub-populations are indicated below each cell subtype.

**Figure 1-figure supplement 2 – Gating strategy to quantify all cell subtypes and sorting major cell subtypes for mitochondrial phenotyping.**

See the methods section for details of labeling cocktails and cell sorter parameters. An initial run of 2M cells was used to establish the six most abundant cell subtypes (targets) for each participant, followed by FACS to obtain at least one 5M aliquot of each target cell subtype. Up to five 5M aliquots were collected per cell subtype, per participant, to establish technical variability in downstream assays.

**Figure 1- figure supplement 3 – Sex differences and age correlations with leukocyte abundance measured by complete blood count.**

(**a**) Forest plot illustrating the effect size (g) of the sex differences in cell proportion derived from the complete blood count (CBC) results (n=21). Fold changes in the raw values are also shown. P-values from non-parametric Mann-Whitney T test. Error bars reflect the 95% C.I. on the effect size. (**b**) Correlation (Spearman’s r) between age and cell proportion derived from the complete blood count. n=21, p<0.05*.

**Figure 2-figure supplement 1 – Associations between CBC cell proportions with mitochondrial features measured in PBMCs.**

Correlations between CBC-based cell type abundance (% of total leukocytes) and PBMCs mitochondrial features for the cohort (n=20).

**Figure 4-figure supplement 1 – Mitochondrial health index (MHI) and coherence of mitochondrial features across cell subtypes.** (**a**) Schematic of the MHI equation reflecting respiratory chain function as the numerator, and markers of mitochondrial content as the denominator, yielding a metric of energy production capacity on a per-mitochondrion basis. (**b**) MHI across immune cell subtypes. Dashed lines are median (thick), with 25th and 75th quartiles (thin). P-values from One-Way non-parametric ANOVA Kruskal-Wallis test with Dunn’s multiple comparison test of subtypes relative to PBMCs, n=12-18 per cell subtype. (**c**) Correlation matrices (Spearman’s r) showing the association between cell subtypes in mitochondrial features. Correlations were not computed for cell subtype pairs with fewer than n=6 observations (gray cell). (**d**) Average effect sizes reflecting the within-person coherence of mitochondrial features across cell types (calculated using Fisher z-transformation). p<0.05*, p<0.001***.

**Figure 4-figure supplement 2 – Associations between subtype-specific enzymatic activities with mitochondrial features measured in PBMCs.**

Correlations of the mitochondrial features measured in each cell subtype and the same mitochondrial feature measured in PBMCs for the cohort (n=12-20). Heatmaps for the cohort show to what extent PBMCs-based measures reflect activities in various immunologically-defined cell subtypes. This data integrates data presented in Figure 4-figure supplement 1, here focused on PBMCs.

**Figure 6-figure supplement 1 – Within-person variability of cell subtype proportions overtime.** (**a**) Within-person variation of cell type proportions across 9 weeks. The FACS-derived raw cell proportions (% of total cells) are shown on the left. Root mean square of successive differences (rMSSD) illustrating the magnitude of variability between successive weeks, and C.V. illustrating the magnitude of variability across the total 9 weeks are provided on the right for each cell subtype, by mitochondrial feature. (**b**) Same as a, but based on CBC results.

**Figure 6-figure supplement 2 – Associations between subtype-specific and CBC cell proportions, and subtype-specific enzymatic activities with mitochondrial features measured in PBMCs.** (**a**) Pairwise correlations of cell subtype proportions obtained from cell sorting with mitochondrial features measured in PBMCs for the repeat participant (n=1 x 9 timepoints). Aggregate correlations are shown as a heatmap (top) and individual scatterplots (bottom). (**b**) Frequency distributions of the effect sizes between PBMC mitochondrial features and cell subtypes proportions for the cohort (Figure 2) and the repeat participant (total correlation pairs=72, for both). (**c**) Correlations between CBC-based cell type abundance (% of total leukocytes) and PBMCs mitochondrial features for the repeat participant (n=1 x 9 timepoints). (**d**) Frequency distribution of effect sizes. (**e**) Correlations of the mitochondrial features measured in each cell subtype and the same mitochondrial feature measured in PBMCs for the repeat participant. Heatmaps for the repeat participant (n=1 x 9 timepoints) show to what extent PBMCs-based measures reflect activities in various immunologically-defined cell subtypes. This data integrates data presented in Figure 6, here focused on PBMCs. (**f**) Frequency distributions of effect sizes for association between PBMC and cell subtype mitochondrial features for the cohort and the repeat participant (total correlation pairs=36, for both), showing a predominance of positive correlations.

**Figure 6-figure supplement 3 – Variability of mitochondrial features across cell subtypes between the cohort and the repeat participant.**

(**a-e**) Side-by-side comparison of mitochondrial features between the cohort (n=12-18 per cell type) and the repeat participant (n=1 x 9 timepoints) across cell subtypes. This figure shows the same data as in main Figure 6h, but for all mitochondrial features. (**f**) Summary of a-e illustrated by a bar graph showing observed variation (C.V.) of mitochondrial features between the cohort and the repeat participant. The technical variation established on a subset of samples and likely represents a conservative overestimation of noise is shown by red lines.

**Figure 7-figure supplement 1 – Operationalization and categorization of mitotypes.**

Chart illustrating mitotype ratios and their simple interpretations.

## Supplementary Files Figure Legends

**Supplementary file 1 – Leukocyte subtypes included in the study.**

Immune cell subtypes included in this study, including a brief summary of their functions and cell surface markers used for immunolabeling and FACS.

---

**Supplementary file 2 – CBC- and FACS-based cell proportions for all study participants and time points.**

Participants are ordered by age. CBC measurements were performed using a Sysmex XN-9000™ instrument, and FACS-based cell proportions were determined using a BD™ Influx cell sorter. (See Appendix 1 for details).

**Supplementary file 3 – Technical variation for each mitochondrial assay and calculated MHI by cell type.**

Coefficients of variation (C.V.s) across 2-5 biological replicates (different 5M cell pellet isolated from the same blood draw) for each cell subtype and PBMCs (See Appendix 1 for details).

**Supplementary file 4 – Recipes for antibody cocktails used to detect cell surface markers for FACS-based cell proportions and sorting.**

(See Appendix 1 for details).

## References

Ackermann, K., Revell, V. L., Lao, O., Rombouts, E. J., Skene, D. J., & Kayser, M. (2012). Diurnal rhythms in blood cell populations and the effect of acute sleep deprivation in healthy young men. Sleep, 35(7), 933–940. doi:10.5665/sleep.1954

Artyomov, M. N., & Van den Bossche, J. (2020). Immunometabolism in the Single-Cell Era. Cell Metabolism. doi:10.1016/j.cmet.2020.09.013

Banas, B., Kost, B. P., & Goebel, F. D. (2004). Platelets, a typical source of error in real-time PCR quantification of mitochondrial DNA content in human peripheral blood cells. Eur J Med Res, 9(8), 371–377.

Beis, D., von Känel, R., Heimgartner, N., Zuccarella-Hackl, C., Bürkle, A., Ehlert, U., & Wirtz, P. H. (2018). The Role of Norepinephrine and α-Adrenergic Receptors in Acute Stress-Induced Changes in Granulocytes and Monocytes. Psychosom Med, 80(7), 649–658. doi:10.1097/psy.0000000000000620

Bektas, A., Schurman, S. H., Gonzalez-Freire, M., Dunn, C. A., Singh, A. K., Macian, F., … Ferrucci, L. (2019). Age-associated changes in human CD4(+) T cells point to mitochondrial dysfunction consequent to impaired autophagy. Aging (Albany NY), 11(21), 9234–9263. doi:10.18632/aging.102438

Bénard, G., Massa, F., Puente, N., Lourenço, J., Bellocchio, L., Soria-Gómez, E., … Marsicano, G. (2012). Mitochondrial CB₁ receptors regulate neuronal energy metabolism. Nat Neurosci, 15(4), 558–564. doi:10.1038/nn.3053

Biino, G., Santimone, I., Minelli, C., Sorice, R., Frongia, B., Traglia, M., … Balduini, C. L. (2013). Age- and sex-related variations in platelet count in Italy: a proposal of reference ranges based on 40987 subjects’ data. PLOS ONE, 8(1), e54289–e54289. doi:10.1371/journal.pone.0054289

Brand, K. (1985). Glutamine and glucose metabolism during thymocyte proliferation. Pathways of glutamine and glutamate metabolism. The Biochemical journal, 228(2), 353–361. doi:10.1042/bj2280353

Brand, M. D., Chen, C. H., & Lehninger, A. L. (1976). Stoichiometry of H+ ejection during respiration-dependent accumulation of Ca2+ by rat liver mitochondria. Journal of Biological Chemistry, 251(4), 968–974. doi:10.1016/S0021-9258(17)33787-0

Butler, L. M., Metson-Scott, T., Felix, J., Abhyankar, A., Rainger, G. E., Farndale, R. W., … Nash, G. B. (2007). Sequential adhesion of platelets and leukocytes from flowing whole blood onto a collagen-coated surface: requirement for a GpVI-binding site in collagen. Thromb Haemost, 97(5), 814–821.

Chacko, B. K., Kramer, P. A., Ravi, S., Johnson, M. S., Hardy, R. W., Ballinger, S. W., & Darley-Usmar, V. M. (2013). Methods for defining distinct bioenergetic profiles in platelets, lymphocytes, monocytes, and neutrophils, and the oxidative burst from human blood. Laboratory investigation; a journal of technical methods and pathology, 93(6), 690–700. doi:10.1038/labinvest.2013.53

Dhabhar, F. S., Malarkey, W. B., Neri, E., & McEwen, B. S. (2012). Stress-induced redistribution of immune cells--from barracks to boulevards to battlefields: a tale of three hormones--Curt Richter Award winner. Psychoneuroendocrinology, 37(9), 1345–1368. doi:10.1016/j.psyneuen.2012.05.008

Dhabhar, F. S., Miller, A. H., Stein, M., McEwen, B. S., & Spencer, R. L. (1994). Diurnal and Acute Stress-Induced Changes in Distribution of Peripheral Blood Leukocyte Subpopulations. Brain, Behavior, and Immunity, 8(1), 66–79. doi:10.1006/brbi.1994.1006

Dixon, N., Li, T., Marion, B., Faust, D., Dozier, S., Molina, A., … Bonkovsky, H. L. (2019). Pilot study of mitochondrial bioenergetics in subjects with acute porphyrias. Mol Genet Metab, 128(3), 228–235. doi:10.1016/j.ymgme.2019.05.010

Du, J., Wang, Y., Hunter, R., Wei, Y., Blumenthal, R., Falke, C., … Manji, H. K. (2009). Dynamic regulation of mitochondrial function by glucocorticoids. Proceedings of the National Academy of Sciences, 106(9), 3543–3548. doi:10.1073/pnas.0812671106

Ehinger, J. K., Morota, S., Hansson, M. J., Paul, G., & Elmér, E. (2016). Mitochondrial Respiratory Function in Peripheral Blood Cells from Huntington’s Disease Patients. Movement disorders clinical practice, 3(5), 472–482. doi:10.1002/mdc3.12308

Fausto-Sterling, A. (2005). The Bare Bones of Sex: Part 1--Sex and Gender. Signs, 30(2), 1491–1527. doi:10.1086/424932

Fisher, A. J., Medaglia, J. D., & Jeronimus, B. F. (2018). Lack of group-to-individual generalizability is a threat to human subjects research. Proceedings of the National Academy of Sciences, 115(27), E6106. doi:10.1073/pnas.1711978115

Gan, Z., Fu, T., Kelly, D. P., & Vega, R. B. (2018). Skeletal muscle mitochondrial remodeling in exercise and diseases. Cell Research, 28(10), 969–980. doi:10.1038/s41422-018-0078-7

Giordano, C., Iommarini, L., Giordano, L., Maresca, A., Pisano, A., Valentino, M. L., … Carelli, V. (2014). Efficient mitochondrial biogenesis drives incomplete penetrance in Leber’s hereditary optic neuropathy. Brain : a journal of neurology, 137(Pt 2), 335–353. doi:10.1093/brain/awt343

Hebert, S. L., Marquet-de Rougé, P., Lanza, I. R., McCrady-Spitzer, S. K., Levine, J. A., Middha, S., … Nair, K. S. (2015). Mitochondrial Aging and Physical Decline: Insights From Three Generations of Women. The journals of gerontology. Series A, Biological sciences and medical sciences, 70(11), 1409–1417. doi:10.1093/gerona/glv086

Hernansanz-Agustín, P., & Enríquez, J. A. (2021). Generation of Reactive Oxygen Species by Mitochondria. Antioxidants (Basel, Switzerland), 10(3), 415. doi:10.3390/antiox10030415

Hurtado-Roca, Y., Ledesma, M., Gonzalez-Lazaro, M., Moreno-Loshuertos, R., Fernandez-Silva, P., Enriquez, J. A., & Laclaustra, M. (2016). Adjusting MtDNA Quantification in Whole Blood for Peripheral Blood Platelet and Leukocyte Counts. PLOS ONE, 11(10), e0163770. doi:10.1371/journal.pone.0163770

Iershov, A., Nemazanyy, I., Alkhoury, C., Girard, M., Barth, E., Cagnard, N., … Panasyuk, G. (2019). The class 3 PI3K coordinates autophagy and mitochondrial lipid catabolism by controlling nuclear receptor PPARα. Nature Communications, 10(1), 1566. doi:10.1038/s41467-019-09598-9

Jang, J. Y., Blum, A., Liu, J., & Finkel, T. (2018). The role of mitochondria in aging. The Journal of Clinical Investigation, 128(9), 3662–3670. doi:10.1172/JCI120842

Johnson, J. L., & Repta, R. (2012). Sex and Gender: Beyond the Binaries.

Jones, N., Vincent, E. E., Cronin, J. G., Panetti, S., Chambers, M., Holm, S. R., … Thornton, C. A. (2019). Akt and STAT5 mediate naïve human CD4+ T-cell early metabolic response to TCR stimulation. Nature Communications, 10(1), 2042. doi:10.1038/s41467-019-10023-4

Karabatsiakis, A., Böck, C., Salinas-Manrique, J., Kolassa, S., Calzia, E., Dietrich, D. E., & Kolassa, I. T. (2014). Mitochondrial respiration in peripheral blood mononuclear cells correlates with depressive subsymptoms and severity of major depression. Translational Psychiatry, 4(6), e397–e397. doi:10.1038/tp.2014.44

Karan, K. R., Trumpff, C., McGill, M. A., Thomas, J. E., Sturm, G., Lauriola, V., … Picard, M. (2020). Mitochondrial respiratory capacity modulates LPS-induced inflammatory signatures in human blood. Brain, Behavior, & Immunity - Health, 5, 100080. doi:10.1016/j.bbih.2020.100080

Kramer, P. A., Ravi, S., Chacko, B., Johnson, M. S., & Darley-Usmar, V. M. (2014). A review of the mitochondrial and glycolytic metabolism in human platelets and leukocytes: implications for their use as bioenergetic biomarkers. Redox biology, 2, 206–210. doi:10.1016/j.redox.2013.12.026

Larsen, S., Nielsen, J., Hansen, C. N., Nielsen, L. B., Wibrand, F., Stride, N., … Hey-Mogensen, M. (2012). Biomarkers of mitochondrial content in skeletal muscle of healthy young human subjects. The Journal of physiology, 590(14), 3349–3360. doi:10.1113/jphysiol.2012.230185

Lee, J. W., Su, Y., Baloni, P., Chen, D., Pavlovitch-Bedzyk, A. J., Yuan, D., … Heath, J. R. (2021). Integrated analysis of plasma and single immune cells uncovers metabolic changes in individuals with COVID-19. Nature Biotechnology. doi:10.1038/s41587-021-01020-4

Lindquist, C., Bjørndal, B., Rossmann, C. R., Svardal, A., Hallström, S., & Berge, R. K. (2018). A fatty acid analogue targeting mitochondria exerts a plasma triacylglycerol lowering effect in rats with impaired carnitine biosynthesis. PLOS ONE, 13(3), e0194978. doi:10.1371/journal.pone.0194978

Maianski, N. A., Geissler, J., Srinivasula, S. M., Alnemri, E. S., Roos, D., & Kuijpers, T. W. (2004). Functional characterization of mitochondria in neutrophils: a role restricted to apoptosis. Cell Death Differ, 11(2), 143–153. doi:10.1038/sj.cdd.4401320

Márquez, E. J., Chung, C.-h., Marches, R., Rossi, R. J., Nehar-Belaid, D., Eroglu, A., … Ucar, D. (2020). Sexual-dimorphism in human immune system aging. Nature Communications, 11(1), 751. doi:10.1038/s41467-020-14396-9

Martínez-Reyes, I., Diebold, L. P., Kong, H., Schieber, M., Huang, H., Hensley, C. T., … Chandel, N. S. (2016). TCA Cycle and Mitochondrial Membrane Potential Are Necessary for Diverse Biological Functions. Mol Cell, 61(2), 199–209. doi:10.1016/j.molcel.2015.12.002

McLaughlin, K. L., Hagen, J. T., Coalson, H. S., Nelson, M. A. M., Kew, K. A., Wooten, A. R., & Fisher-Wellman, K. H. (2020). Novel approach to quantify mitochondrial content and intrinsic bioenergetic efficiency across organs. Scientific Reports, 10(1), 17599. doi:10.1038/s41598-020-74718-1

Mengel-From, J., Thinggaard, M., Dalgård, C., Kyvik, K. O., Christensen, K., & Christiansen, L. (2014). Mitochondrial DNA copy number in peripheral blood cells declines with age and is associated with general health among elderly. Hum Genet, 133(9), 1149–1159. doi:10.1007/s00439-014-1458-9

Michalek, R. D., Gerriets, V. A., Jacobs, S. R., Macintyre, A. N., MacIver, N. J., Mason, E. F., … Rathmell, J. C. (2011). Cutting edge: distinct glycolytic and lipid oxidative metabolic programs are essential for effector and regulatory CD4+ T cell subsets. Journal of immunology (Baltimore, Md. : 1950), 186(6), 3299–3303. doi:10.4049/jimmunol.1003613

Moore, A. Z., Ding, J., Tuke, M. A., Wood, A. R., Bandinelli, S., Frayling, T. M., & Ferrucci, L. (2018). Influence of cell distribution and diabetes status on the association between mitochondrial DNA copy number and aging phenotypes in the InCHIANTI study. Aging cell, 17(1), e12683. doi:10.1111/acel.12683

Nicholls, D. G., & Ferguson, S. J. (2013). Bioenergetics: Academic Press.

Nicoli, F., Papagno, L., Frere, J. J., Cabral-Piccin, M. P., Clave, E., Gostick, E., … Appay, V. (2018). Naïve CD8+ T-Cells Engage a Versatile Metabolic Program Upon Activation in Humans and Differ Energetically From Memory CD8+ T-Cells. Frontiers in Immunology, 9(2736). doi:10.3389/fimmu.2018.02736

Nikolich-Žugich, J. (2014). Aging of the T cell compartment in mice and humans: from no naive expectations to foggy memories. Journal of immunology (Baltimore, Md. : 1950), 193(6), 2622–2629. doi:10.4049/jimmunol.1401174

Nomura, M., Liu, J., Rovira, I. I., Gonzalez-Hurtado, E., Lee, J., Wolfgang, M. J., & Finkel, T. (2016). Fatty acid oxidation in macrophage polarization. Nature Immunology, 17(3), 216–217. doi:10.1038/ni.3366

Patin, E., Hasan, M., Bergstedt, J., Rouilly, V., Libri, V., Urrutia, A., … The Milieu Intérieur, C. (2018). Natural variation in the parameters of innate immune cells is preferentially driven by genetic factors. Nature Immunology, 19(3), 302–314. doi:10.1038/s41590-018-0049-7

Pearce, E. L., Poffenberger, M. C., Chang, C.-H., & Jones, R. G. (2013). Fueling Immunity: Insights into Metabolism and Lymphocyte Function. Science, 342(6155), 1242454. doi:10.1126/science.1242454

Picard, M. (2021). Blood mitochondrial DNA copy number: What are we counting? Mitochondrion, 60, 1–11. doi:https://doi.org/10.1016/j.mito.2021.06.010

Picard, M., Jung, B., Liang, F., Azuelos, I., Hussain, S., Goldberg, P., … Petrof, B. J. (2012). Mitochondrial dysfunction and lipid accumulation in the human diaphragm during mechanical ventilation. Am J Respir Crit Care Med, 186(11), 1140–1149. doi:10.1164/rccm.201206-0982OC

Picard, M., & McEwen, B. S. (2018). Psychological Stress and Mitochondria: A Systematic Review. Psychosom Med, 80(2), 141–153. doi:10.1097/psy.0000000000000545

Picard, M., Prather, A. A., Puterman, E., Cuillerier, A., Coccia, M., Aschbacher, K., … Epel, E. S. (2018). A Mitochondrial Health Index Sensitive to Mood and Caregiving Stress. Biol Psychiatry, 84(1), 9–17. doi:10.1016/j.biopsych.2018.01.012

Picard, M., Trumpff, C., & Burelle, Y. (2019). Mitochondrial psychobiology: foundations and applications. Current opinion in behavioral sciences, 28, 142–151. doi:10.1016/j.cobeha.2019.04.015

Picard, M., Wallace, D. C., & Burelle, Y. (2016). The rise of mitochondria in medicine. Mitochondrion, 30, 105–116. doi:10.1016/j.mito.2016.07.003

Picard, M., Zhang, J., Hancock, S., Derbeneva, O., Golhar, R., Golik, P., … Wallace, D. C. (2014). Progressive increase in mtDNA 3243A>G heteroplasmy causes abrupt transcriptional reprogramming. Proceedings of the National Academy of Sciences of the United States of America, 111(38), E4033–4042. doi:10.1073/pnas.1414028111

Pyle, A., Burn, D. J., Gordon, C., Swan, C., Chinnery, P. F., & Baudouin, S. V. (2010). Fall in circulating mononuclear cell mitochondrial DNA content in human sepsis. Intensive Care Medicine, 36(6), 956–962. doi:10.1007/s00134-010-1823-7

Ritz, S. A., Antle, D. M., Côté, J., Deroy, K., Fraleigh, N., Messing, K., … Mergler, D. (2014). First steps for integrating sex and gender considerations into basic experimental biomedical research. Faseb j, 28(1), 4–13. doi:10.1096/fj.13-233395

Ron-Harel, N., Ghergurovich, J. M., Notarangelo, G., LaFleur, M. W., Tsubosaka, Y., Sharpe, A. H., … Haigis, M. C. (2019). T Cell Activation Depends on Extracellular Alanine. Cell Reports, 28(12), 3011–3021.e3014. doi:10.1016/j.celrep.2019.08.034

Segerstrom, S. C., Sephton, S. E., & Westgate, P. M. (2017). Intraindividual variability in cortisol: Approaches, illustrations, and recommendations. Psychoneuroendocrinology, 78, 114–124. doi:10.1016/j.psyneuen.2017.01.026

Shim, H. B., Arshad, O., Gadawska, I., Côté, H. C. F., & Hsieh, A. Y. Y. (2020). Platelet mtDNA content and leukocyte count influence whole blood mtDNA content. Mitochondrion, 52, 108–114. doi:10.1016/j.mito.2020.03.001

Short, K. R., Bigelow, M. L., Kahl, J., Singh, R., Coenen-Schimke, J., Raghavakaimal, S., & Nair, K. S. (2005). Decline in skeletal muscle mitochondrial function with aging in humans. Proceedings of the National Academy of Sciences of the United States of America, 102(15), 5618–5623. doi:10.1073/pnas.0501559102

Silaidos, C., Pilatus, U., Grewal, R., Matura, S., Lienerth, B., Pantel, J., & Eckert, G. P. (2018). Sex-associated differences in mitochondrial function in human peripheral blood mononuclear cells (PBMCs) and brain. Biology of Sex Differences, 9(1), 34. doi:10.1186/s13293-018-0193-7

Turner, N., Bruce, C. R., Beale, S. M., Hoehn, K. L., So, T., Rolph, M. S., & Cooney, G. J. (2007). Excess lipid availability increases mitochondrial fatty acid oxidative capacity in muscle: evidence against a role for reduced fatty acid oxidation in lipid-induced insulin resistance in rodents. Diabetes, 56(8), 2085–2092. doi:10.2337/db07-0093

Tyrrell, D. J., Bharadwaj, M. S., Van Horn, C. G., Marsh, A. P., Nicklas, B. J., & Molina, A. J. (2015). Blood-cell bioenergetics are associated with physical function and inflammation in overweight/obese older adults. Exp Gerontol, 70, 84–91. doi:10.1016/j.exger.2015.07.015

Urata, M., Koga-Wada, Y., Kayamori, Y., & Kang, D. (2008). Platelet contamination causes large variation as well as overestimation of mitochondrial DNA content of peripheral blood mononuclear cells. Ann Clin Biochem, 45(Pt 5), 513–514. doi:10.1258/acb.2008.008008

van der Windt, G. J., Everts, B., Chang, C. H., Curtis, J. D., Freitas, T. C., Amiel, E., … Pearce, E. L. (2012). Mitochondrial respiratory capacity is a critical regulator of CD8+ T cell memory development. Immunity, 36(1), 68–78. doi:10.1016/j.immuni.2011.12.007

Ventura-Clapier, R., Piquereau, J., Veksler, V., & Garnier, A. (2019). Estrogens, Estrogen Receptors Effects on Cardiac and Skeletal Muscle Mitochondria. Frontiers in Endocrinology, 10(557). doi:10.3389/fendo.2019.00557

Verhoeven, J. E., Révész, D., Picard, M., Epel, E. E., Wolkowitz, O. M., Matthews, K. A., … Puterman, E. (2018). Depression, telomeres and mitochondrial DNA: between- and within-person associations from a 10-year longitudinal study. Mol Psychiatry, 23(4), 850–857. doi:10.1038/mp.2017.48

Wachsmuth, M., Hübner, A., Li, M., Madea, B., & Stoneking, M. (2016). Age-Related and Heteroplasmy-Related Variation in Human mtDNA Copy Number. PLOS Genetics, 12(3), e1005939. doi:10.1371/journal.pgen.1005939

Wallace, Douglas C. (2015). Mitochondrial DNA Variation in Human Radiation and Disease. Cell, 163(1), 33–38. doi:10.1016/j.cell.2015.08.067

Weiss, S. L., Selak, M. A., Tuluc, F., Perales Villarroel, J., Nadkarni, V. M., Deutschman, C. S., & Becker, L. B. (2015). Mitochondrial dysfunction in peripheral blood mononuclear cells in pediatric septic shock. Pediatric critical care medicine : a journal of the Society of Critical Care Medicine and the World Federation of Pediatric Intensive and Critical Care Societies, 16(1), e4–e12. doi:10.1097/PCC.0000000000000277

Ye, K., Lu, J., Ma, F., Keinan, A., & Gu, Z. (2014). Extensive pathogenicity of mitochondrial heteroplasmy in healthy human individuals. Proceedings of the National Academy of Sciences, 111(29), 10654. doi:10.1073/pnas.1403521111

Yu-Wai-Man, P., Sitarz, K. S., Samuels, D. C., Griffiths, P. G., Reeve, A. K., Bindoff, L. A., … Chinnery, P. F. (2010). OPA1 mutations cause cytochrome c oxidase deficiency due to loss of wild-type mtDNA molecules. Human molecular genetics, 19(15), 3043–3052. doi:10.1093/hmg/ddq209

Zhang, J., Li, M., & He, Y. (2015). Large population study for age˔ and gender- related variations of platelet indices in Southwest China healthy adults. Hematology & Transfusion International Journal, 1.

Zhang, R., Wang, Y., Ye, K., Picard, M., & Gu, Z. (2017). Independent impacts of aging on mitochondrial DNA quantity and quality in humans. BMC Genomics, 18(1), 890. doi:10.1186/s12864-017-4287-0

## References

Chiu, S. and A. Bharat (2016). “Role of monocytes and macrophages in regulating immune response following lung transplantation.” Current opinion in organ transplantation 21(3): 239–245.

Gasper, D. J., M. M. Tejera and M. Suresh (2014). “CD4 T-cell memory generation and maintenance.” Critical reviews in immunology 34(2): 121–146.

Hoffman, W., F. G. Lakkis and G. Chalasani (2016). “B Cells, Antibodies, and More.” Clinical journal of the American Society of Nephrology : CJASN 11(1): 137–154.

Luckheeram, R. V., R. Zhou, A. D. Verma and B. Xia (2012). “CD4⁺T cells: differentiation and functions.” Clinical & developmental immunology 2012: 925135–925135.

Mahnke, Y. D., T. M. Brodie, F. Sallusto, M. Roederer and E. Lugli (2013). “The who’s who of T-cell differentiation: Human memory T-cell subsets.” European Journal of Immunology 43(11): 2797–2809.

Martin, M. D. and V. P. Badovinac (2018). “Defining Memory CD8 T Cell.” Frontiers in Immunology 9(2692).

Mayadas, T. N., X. Cullere and C. A. Lowell (2014). “The Multifaceted Functions of Neutrophils.” Annual Review of Pathology: Mechanisms of Disease 9(1): 181–218.

Stubbe, M., N. Vanderheyde, M. Goldman and A. Marchant (2006). “Antigen-specific central memory CD4+ T lymphocytes produce multiple cytokines and proliferate in vivo in humans.” J Immunol 177(11): 8185–8190.

Tian, Y., M. Babor, J. Lane, V. Schulten, V. S. Patil, G. Seumois, S. L. Rosales, Z. Fu, G. Picarda, J. Burel, J. Zapardiel-Gonzalo, R. N. Tennekoon, A. D. De Silva, S. Premawansa, G. Premawansa, A. Wijewickrama, J. A. Greenbaum, P. Vijayanand, D. Weiskopf, A. Sette and B. Peters (2017). “Unique phenotypes and clonal expansions of human CD4 effector memory T cells re-expressing CD45RA.” Nature Communications 8(1): 1473.

Vivier, E., E. Tomasello, M. Baratin, T. Walzer and S. Ugolini (2008). “Functions of natural killer cells.” Nature Immunology 9: 503–510.

Zhang, N. and M. J. Bevan (2011). “CD8(+) T cells: foot soldiers of the immune system.” Immunity 35(2): 161–168.

