## Supplementary figures and images for "Mitochondrial phenotypes in purified human immune cell subtypes and cell mixtures"

### Figure 1-figure supplement 1

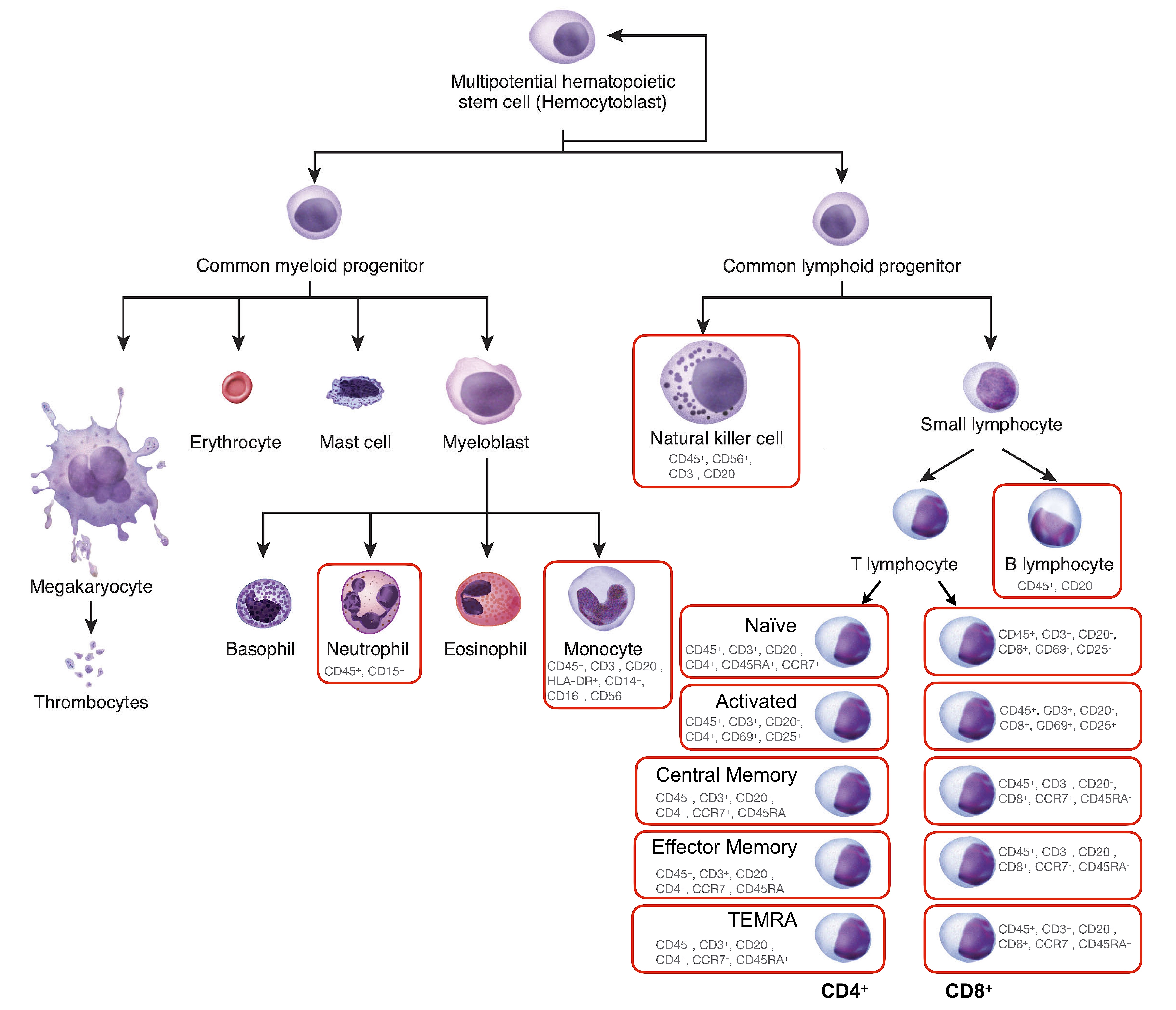

### Figure 1-figure supplement 2

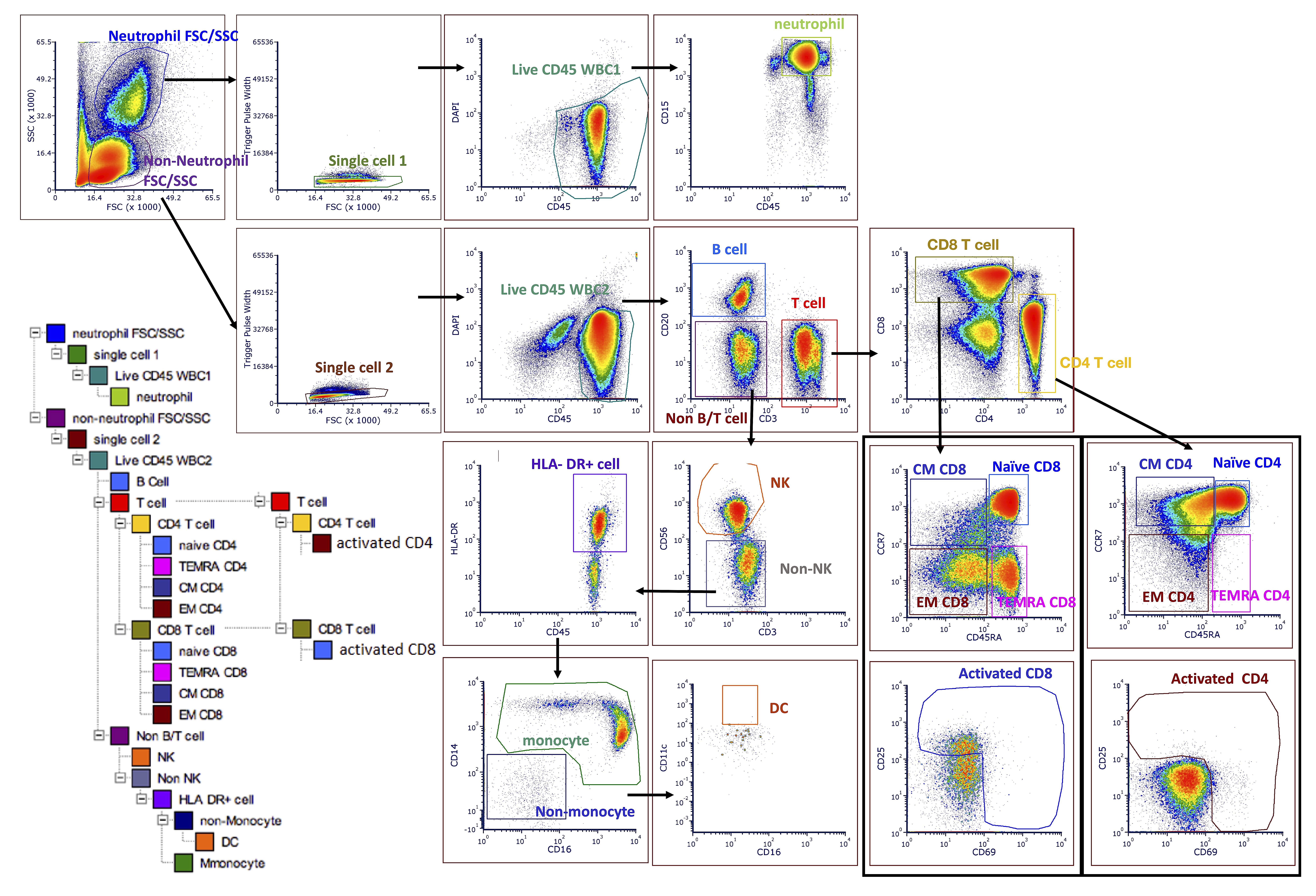

### Figure 1-figure supplement 3

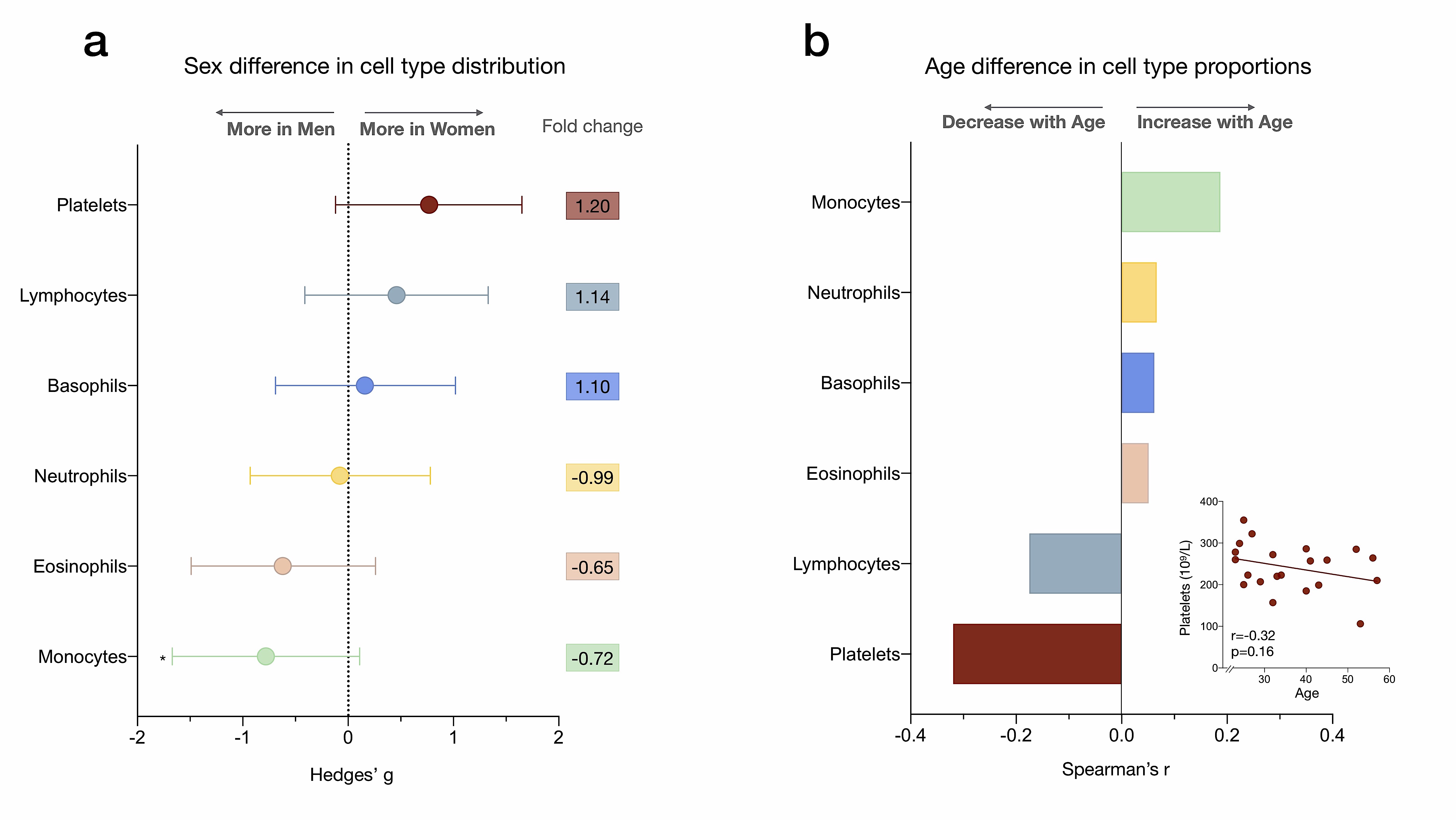

### Figure 2-figure supplement 1

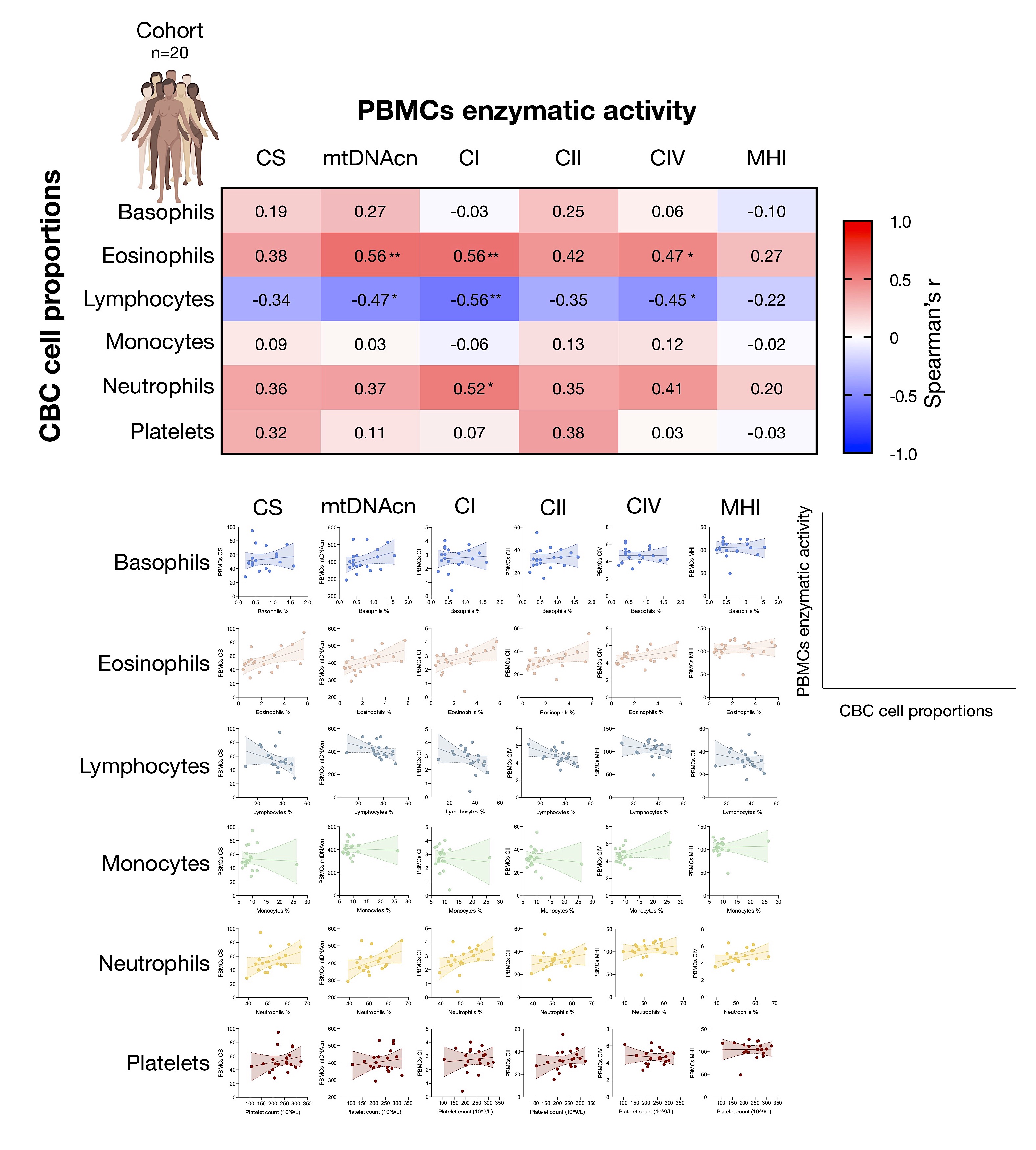

### Figure 4-figure supplement 1

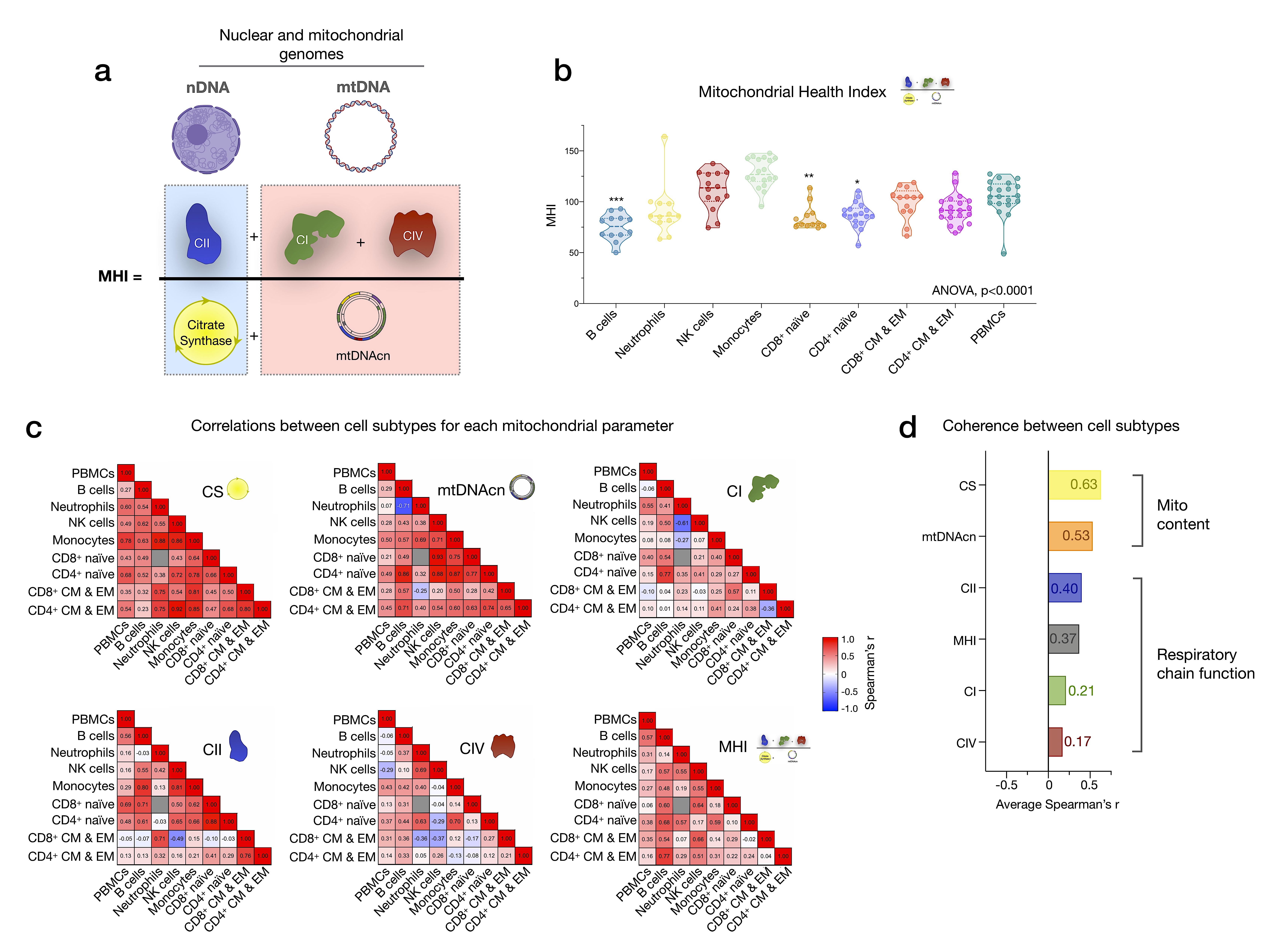

### Figure 4-figure supplement 2

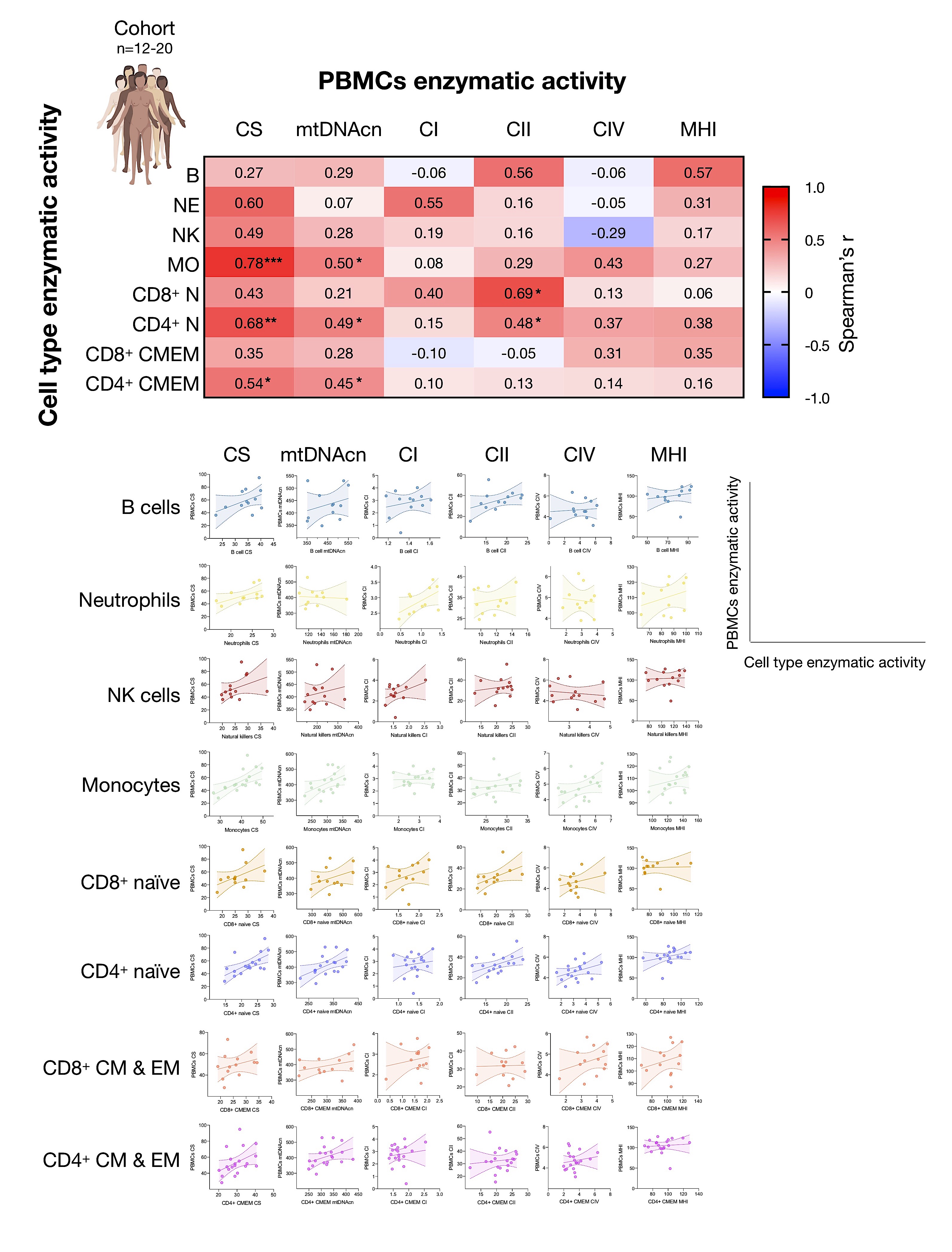

### Figure 6-figure supplement 1

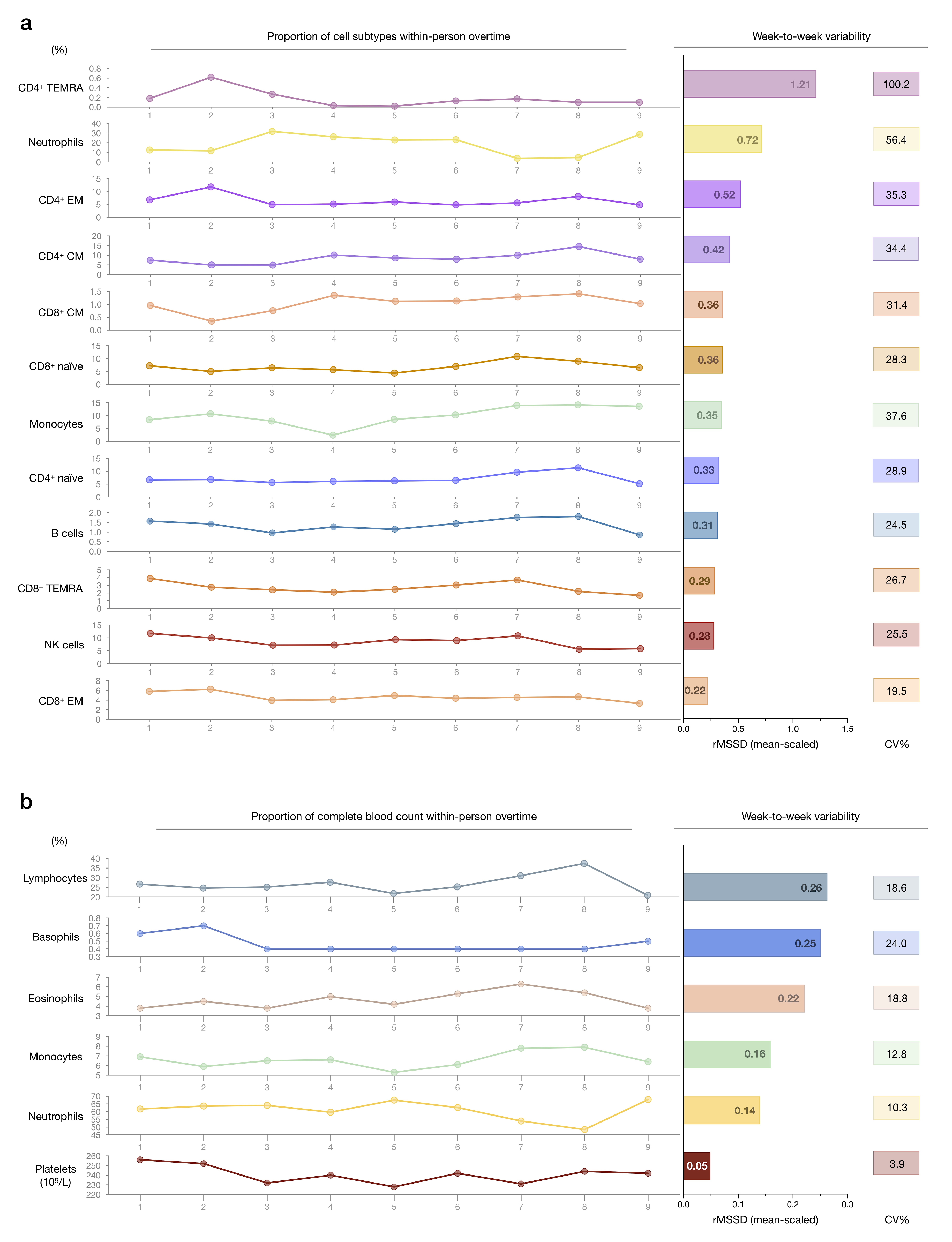

### Figure 6-figure supplement 2

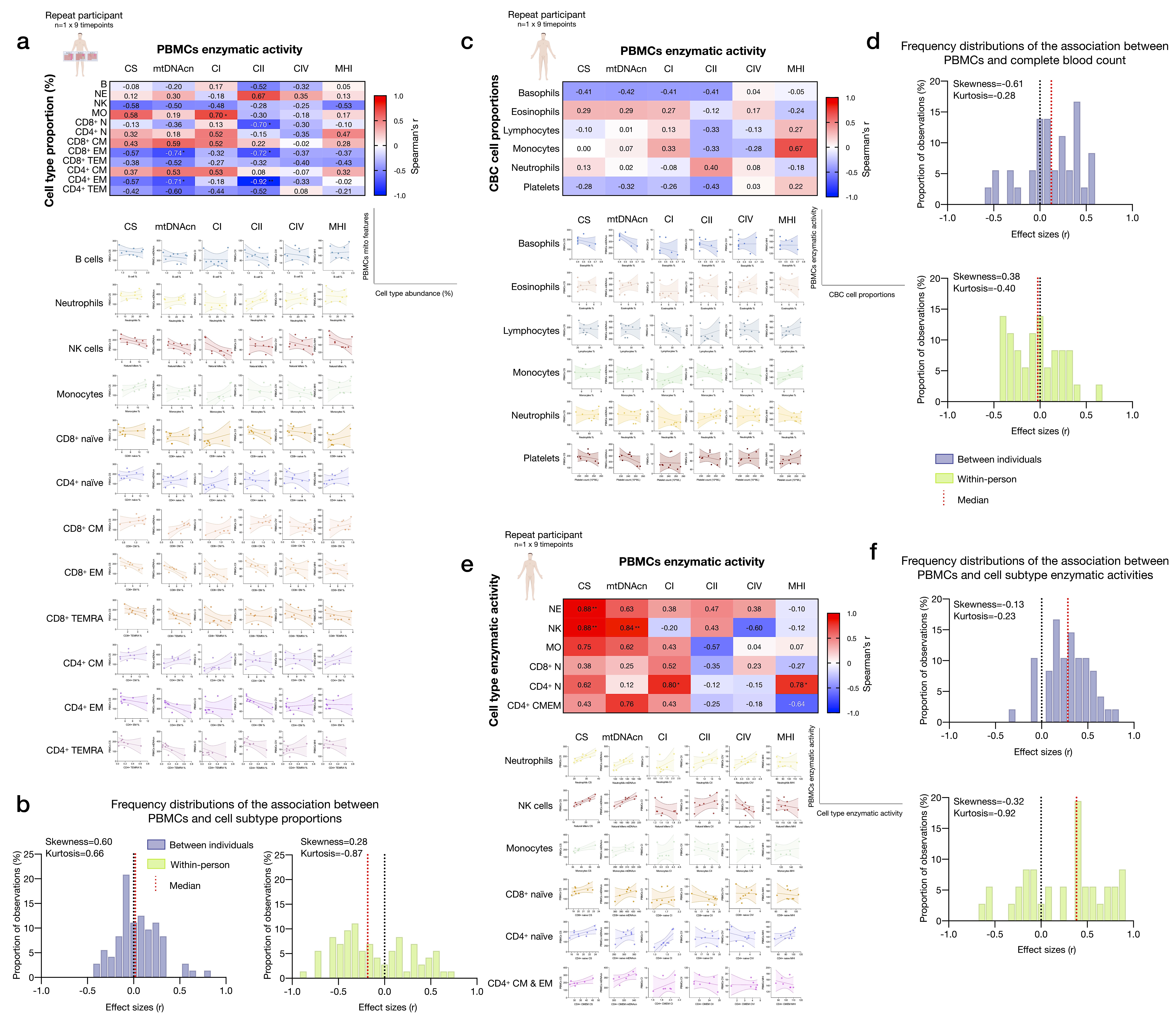

### Figure 6-figure supplement 3

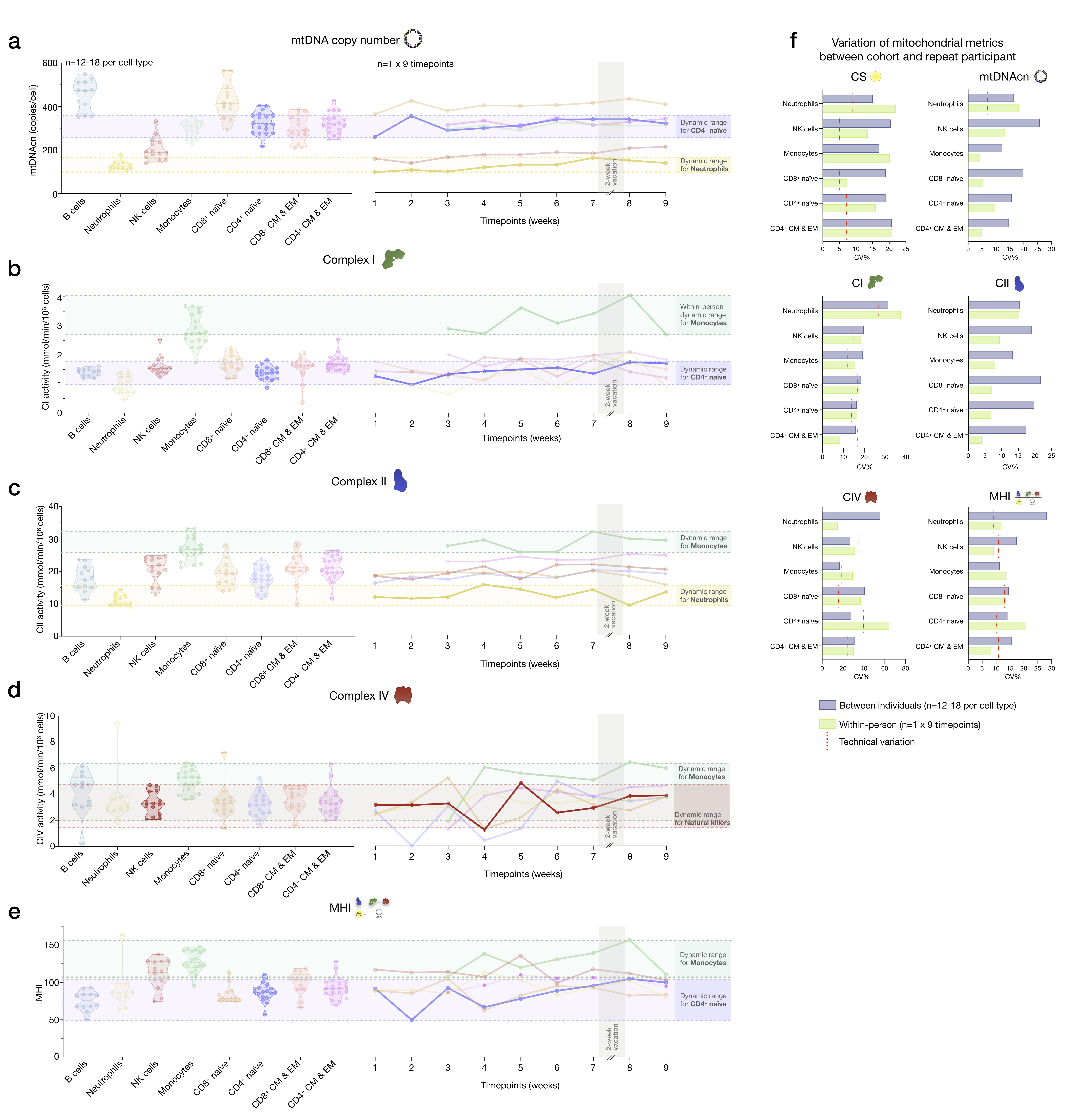

### Figure 7-figure supplement 1

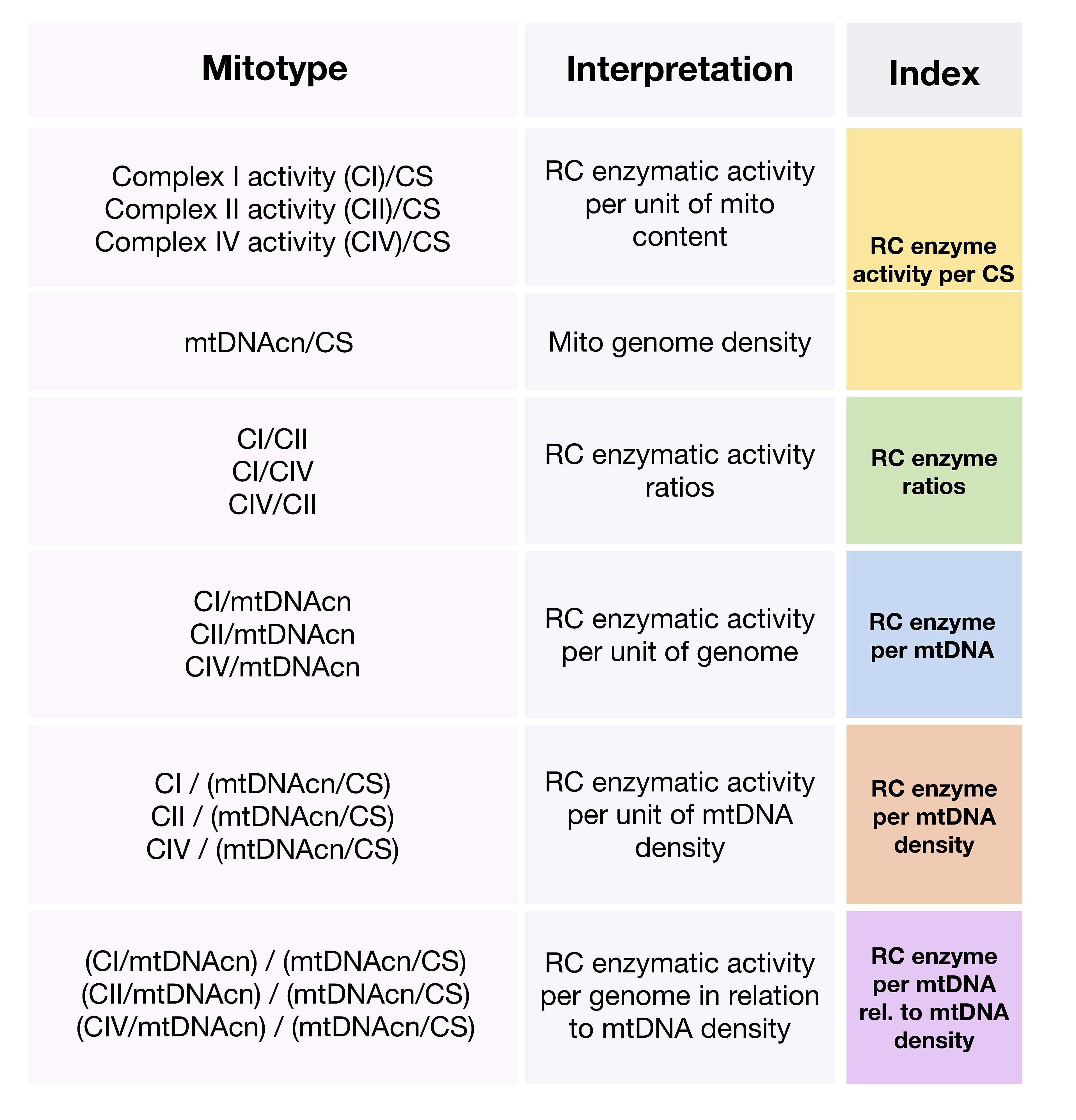

### Supplementary file 1

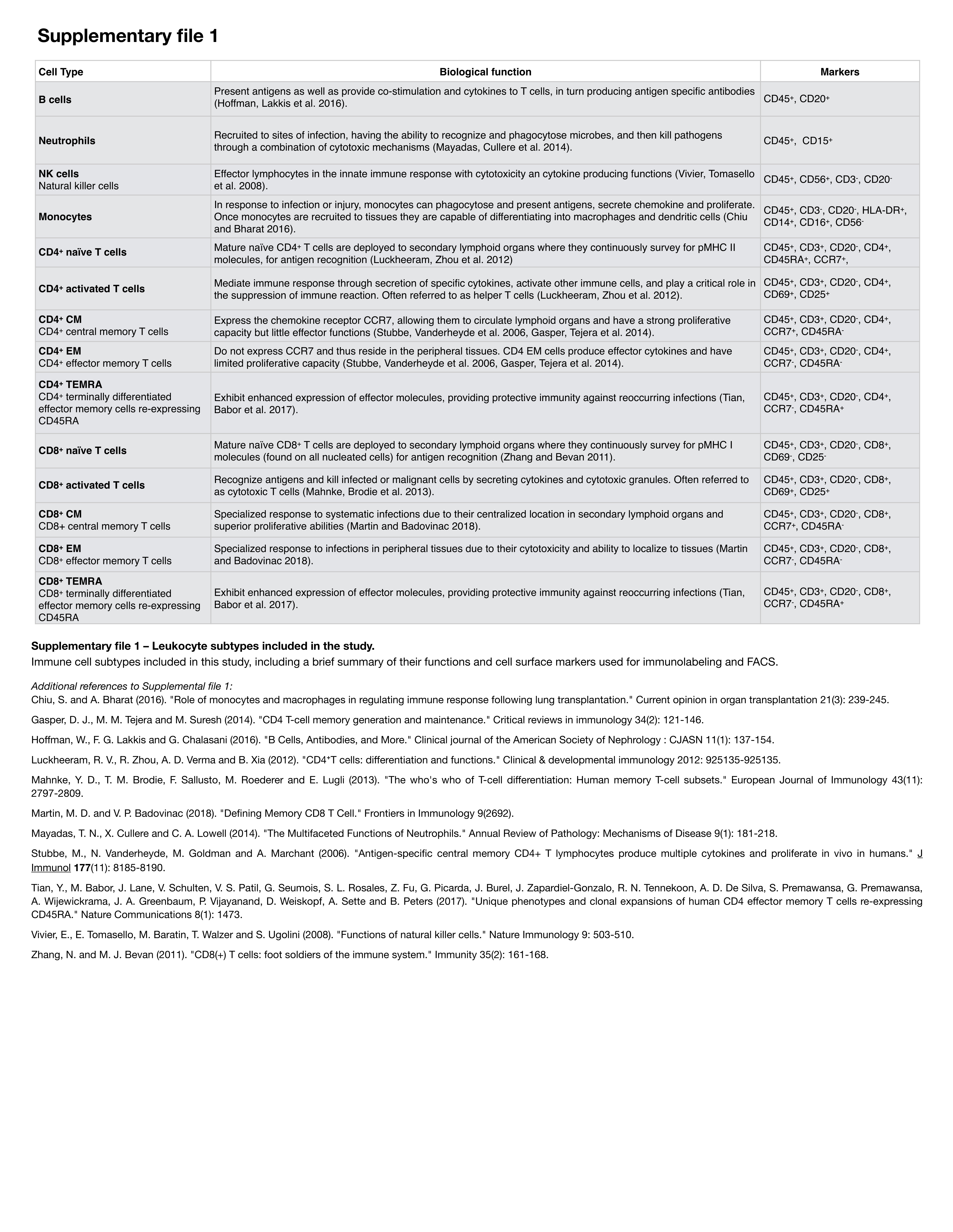

### Supplementary file 2

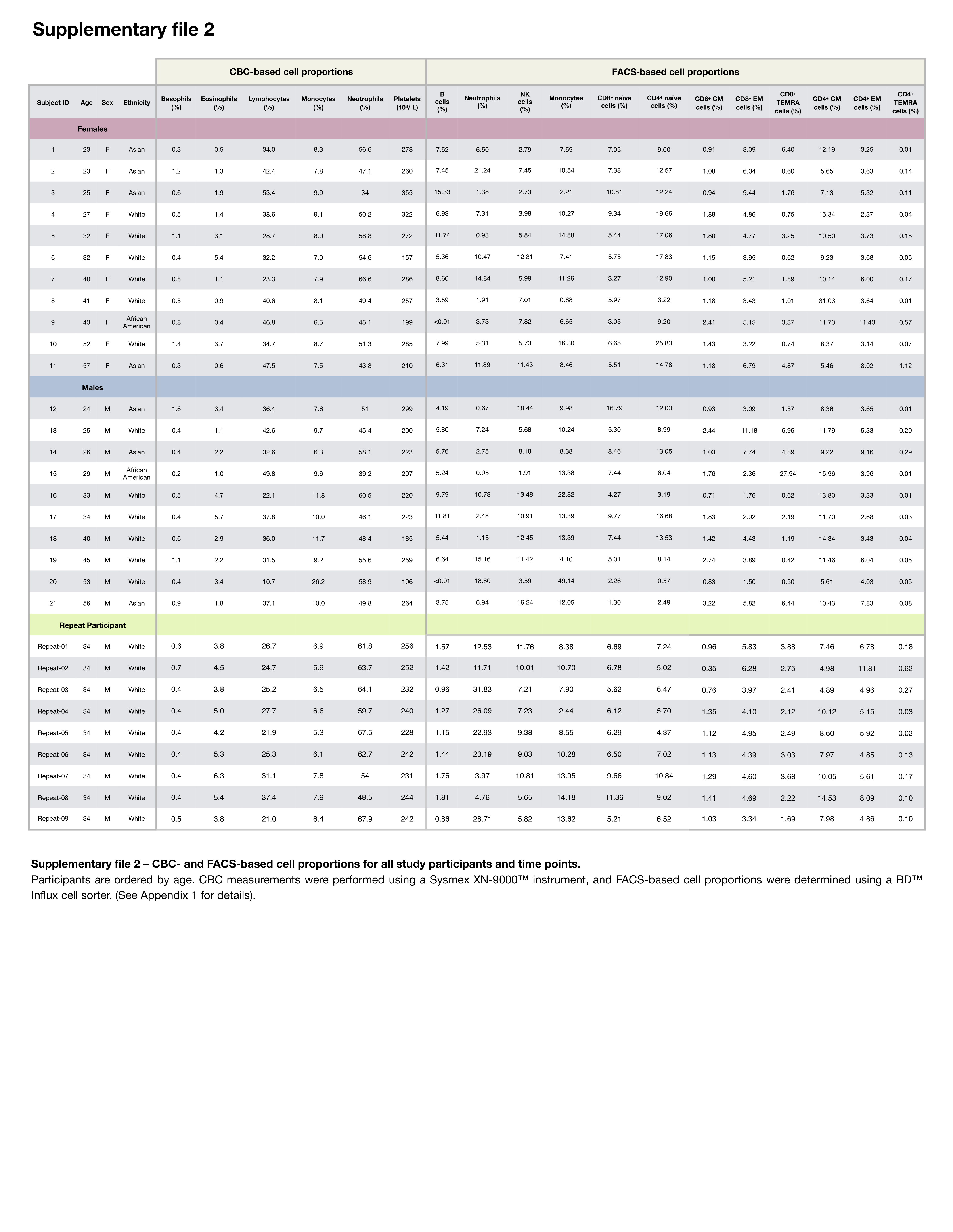

### Supplementary file 3

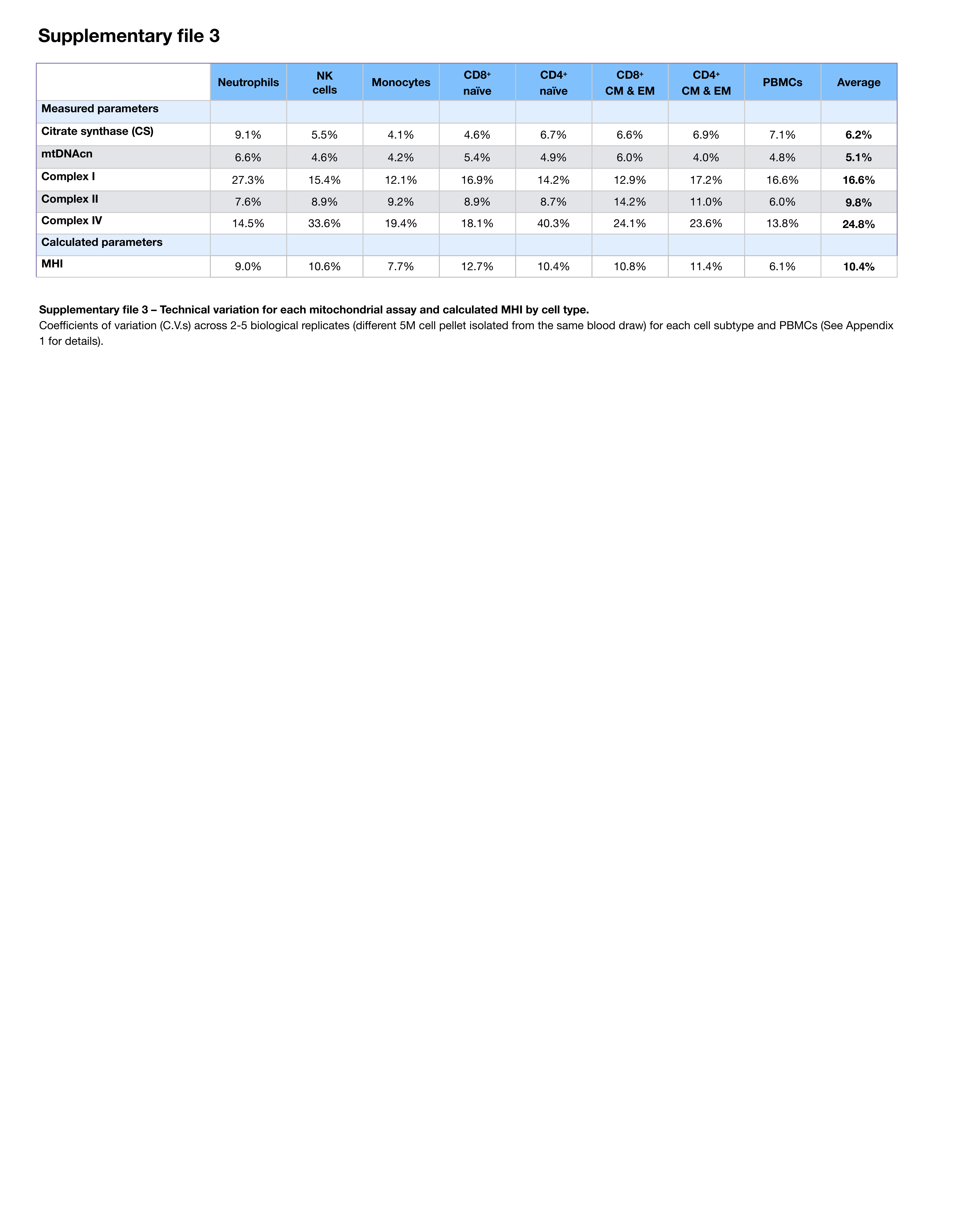

### Supplementary file 4

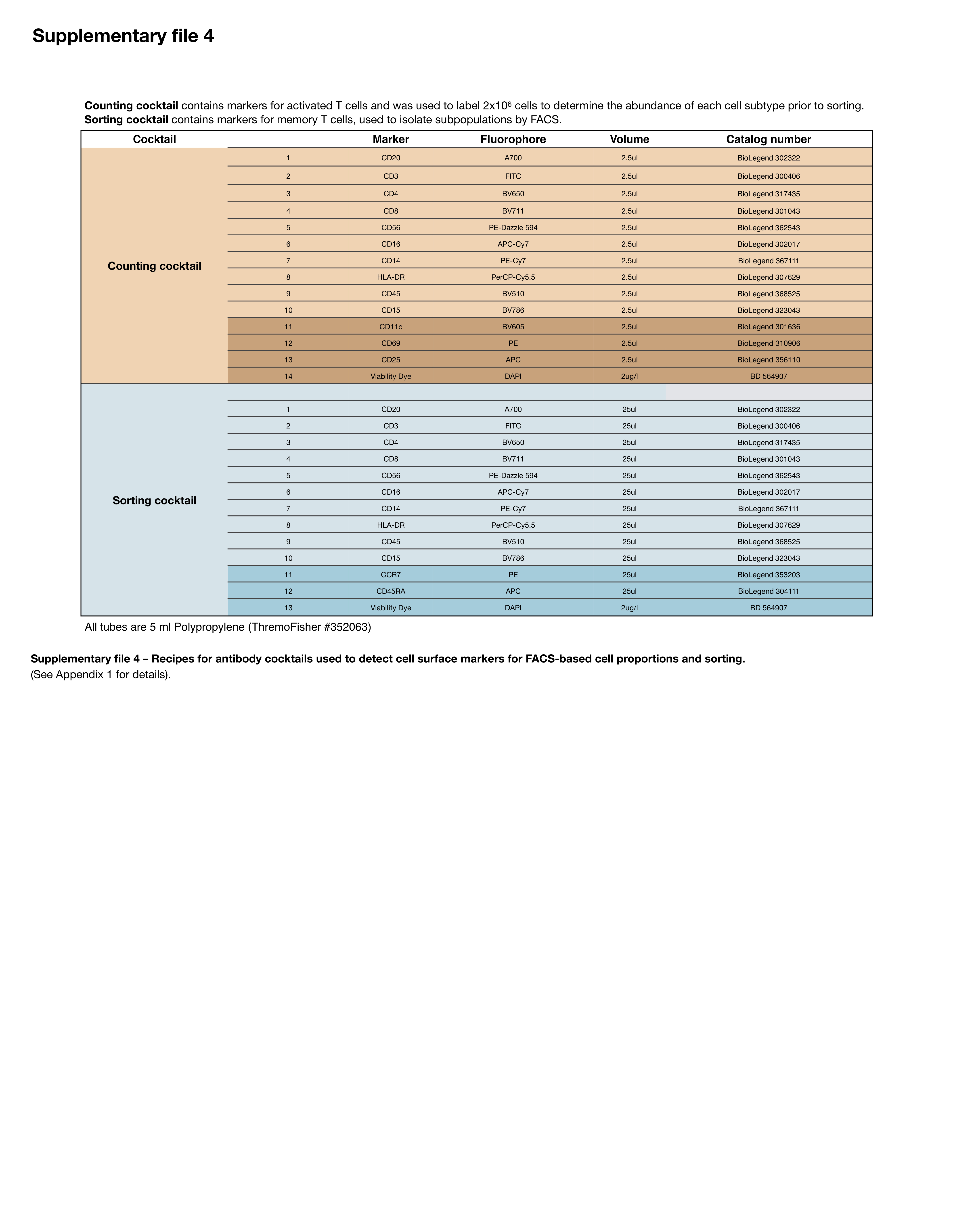
