## Appendix 1 for "Mitochondrial phenotypes in purified human immune cell subtypes and cell mixtures"

### Supplemental Methods and Procedures

*This Appendix is an extended version of the Methods section in the main text. It includes additional technical details to facilitate the implementation and reproducibility of methods used in this study.*

*Blood Collection*

*A total o*f 100 ml of blood was drawn from the antecubital vein for each participant and included one EDTA tube (Cat# BD367841, 2 ml) for complete blood count (CBC), two SST coated tubes (Cat#BD367986, 5 ml) for hormonal measures and blood biochemistry in serum, and 11 Acid Dextrose A (ACD-A) tubes (Cat# BD364606, 8.5 ml) for leukocyte isolation and mitochondrial analyses, in order of collection. All tubes were gently inverted 10-12 times immediately after draw to ensure proper mixing. The EDTA and SST tubes for hematological and blood biochemistry were processed by the Center for Advanced Laboratory Medicine (CALM) Lab, and the ACD-A tubes were further processed for cell isolation.

*PBMCs and leukocyte isolation*

Ficoll 1077 and 1119 (Sigma), Hanks Balanced Salt Sodium (HBSS) without phenol red, calcium and magnesium (Life Technologies, Cat# 14175103) supplemented with 2% BSA (Sigma, Cat# A9576) (HBSS/BSA), HBSS/BSA supplemented with 1 mM of EDTA (Sigma, Cat# E9884) (HBSS/BSA/EDTA), and FBS (Thermofisher, cat# 10437036) were brought to room temperature overnight. PBMCs were isolated on 15 ml of low density Ficoll 1077 in a 50 ml conical tube, and total leukocytes were separated on 15ml of higher density Ficoll 1119 distributed across 7 conical tubes. Blood was first pooled and diluted with HBSS/BSA in a 1:1 ratio, and 25 ml of diluted blood was carefully layered on Ficoll and then centrifuged immediately at 700 x g for 30 minutes (no brake) in a horizontal rotor (swing-out head) tabletop centrifuge, at room temperature. Immediately after centrifugation, cells at the interface were collected and washed in 50 ml HBSS/BSA and centrifuged at 700 x g for 10 minutes. Supernatants were discarded and leukocyte pellets washed again in HBSS/BSA/EDTA and centrifuged at 700 x g for 10 minutes to limit platelet contamination. Low concentration EDTA (1mM) was used to prevent cell-cell adhesion or platelet activation, but a higher concentration was not used to avoid perturbing downstream mitochondrial assays.

To perform cell count, both i) PBMCs (1:10 dilution) and ii) total leukocytes (1:100 dilution) were resuspended in 1 ml of HBSS/BSA/EDTA and counted on the Countess II FL Automated Cell Counter (Thermo Fisher Scientific, Cat# AMQAF1000) in a 1:1 dilution with trypan blue. Counts were performed in duplicates. If the difference between duplicates >10%, the count was repeated and the grand mean of the cell counts taken. Pellets of 5 million PBMCs were aliquoted and frozen at -80°C for mitochondrial assays.

*Immunolabeling of cell surface markers*

Two antibody cocktails meant for i) cell counting (Cocktail 1) and ii) cell sorting (Cocktail 2), were prepared for fluorescence-activated cell sorting (FACS) (see Supplementary file 4 for details). Antibodies were gently mixed, kept at 4°C, and shielded from light to avoid bleaching. Cocktail 1 (containing cell surface markers for activated T lymphocytes) was prepared with 17.5 ul HBSS/BSA/EDTA and 2.5 ul per antibody (13 markers, 32.5 ul total antibody mix), for a total of 50 ul. Cocktail 2 was prepared with 200 ul HBSS/BSA/EDTA and 25 ul per antibody (12 markers, 300 ul total sorting antibody mix), for a total of 500 ul.

Prior to each study visit, cell collection tubes (Cat#: 352063, polypropylene, 5 ml) were coated with 4.5ml of DMEM/10% FBS media to minimize cell-tube adhesion and maximize the recovery of sorted cells. Tubes were incubated for 24 hours at room temperature and stored at 4°C until use, and decanted prior to use. Two coated polypropylene tubes were used for the FACS-ready antibody-labeled leukocytes, and an additional 60 coated polypropylene falcon tubes were decanted and 500 ul of media (DMEM/10% FBS) was added to receive sorted cells.

Prior to immunolabeling, total leukocytes were incubated with blocking buffer (to block non-specific binding to FC receptors) at a 1:10 dilution and incubated at room temperature for 10 minutes. A 2 million cell aliquot was diluted to a final volume of 100 ul with HBSS/BSA/EDTA and combined with 50 ul of Cocktail 1. The remainder of total leukocytes (~100M cells) were incubated with 500 ul of Cocktail 2 for 20 minutes in the dark, at room temperature. Both cell preparations were then washed with 5 ml of HBSS/BSA/EDTA and centrifuged at 700 x g for 5 minutes. Using the propylene tubes, Cocktail 1 cells were resuspended in 200 ul of HBSS/BSA/EDTA, and total leukocytes for FACS were resuspended to final concentration of 20 million cells/ml with HBSS/BSA/EDTA.

*Fluorescence-activated cell sorting (FACS)*

Cells labeled with the Cocktail 1 (counting) panel was only used for data acquisition and phenotype analysis. Cells labeled with the Cocktail 2 (sorting) panel was FACS sorted using a BD™ Influx cell sorter to isolate the subpopulations from peripheral blood. The sorter was controlled using BD FACS Sortware. Cells were sorted using 100 um size nozzle and under the sheath pressure of 20 psi. Sorting speed was kept around 11,000-12,000 events/second. Cell concentration for sorting was measured at about $15x10^6$ cells per ml. Cell sorter drop frequency was 37 KHz, stream focus was 10%, maximum drop charge was 110 Volts. A six-way sorting, 1.0 drop pure sort mode was used to sort the cell subpopulations. Stream deflections were -84, -65, -32, 32, 65, and 84 for six-way sort from left to right. For each participant, 1 million cells (Cocktail 1 panel) were run first to calculate the potential yield of each subpopulation, including neutrophils, B lymphocytes, monocytes, NK cells, naïve $CD4^+$ and $CD8^+$, CM $CD4^+$ and $CD8^+$, EM $CD4^+$ and $CD8^+$, and TEMRA $CD4^+$ and $CD8^+$ lymphocytes (total cell number x percentage of each subpopulation). The six most abundant subpopulations were sorted. Purity checks were performed on all sorted subpopulations to ensure the instrument performance was good enough to reach the sorted population purity >95%. Raw data (.fcs file) was exported for further analysis on FCS Express 7 research version.

*Processing and storage of sorted cells*

Following flow cytometry, sorted cell subtypes were transferred and pooled by pipetting about half of each collection tube (2.5 ml) into larger falcon tubes, gently vortexing to liberate cells that may have adhered to the tube wall, and the remaining volume pipetted into the transfer tube. HBSS/2% BSA was used as necessary to equilibrate and cells were centrifuged at 1,000 x g for 5 minutes. Following centrifugation, each cell pellet was isolated by gently decanting the supernatant and re-suspended into 1ml of HBSS/2% BSA. The resulting purified cell suspensions were transferred to a 1.5 ml Eppendorf tube for each cell type, centrifuged at 2,000 x g for 2 minutes at 4°C, and the supernatant carefully removed to leave a dry cell pellet. Samples were stored in liquid nitrogen for 4-12 months (-170°C) until mitochondrial biochemistry and mtDNAcn analyses were performed as a single batch.

*Mitochondrial enzymatic activities*

Samples were thawed and homogenized in preparation for enzymatic activity measurements with one tungsten bead and 500 ul of homogenization buffer (1 mM EDTA, 50 mM Triethanolamine). Tubes were transferred to a pre-chilled rack and homogenized using a Tissue Lyser (Qiagen cat# 85300) at 30 cycles/second for 1 minute. Samples were then incubated for 5 minutes at 4°C, and homogenization was repeated for 1 minute and the samples were returned to ice ready for enzymatic assays.

Mitochondrial enzyme activities were quantified spectrophotometrically for citrate synthase (CS), cytochrome c oxidase (COX, Complex IV), succinate dehydrogenase (SDH, Complex II), and NADH dehydrogenase (Complex I) as described previously (Picard et al., 2018) with minor modifications. Each sample was measured in triplicates for each enzymatic assay (3 wells for total activity and 3 wells for non-specific activity, except for the COX assay where a single non-specific activity value is determined across 30 wells). Homogenate volumes used for each reaction were: CS: 10 ul, COX and SDH: 20 ul, Complex I: 15 ul.

CS activity was measured by detecting the increase in absorbance at 412 nm, in a reaction buffer (200 mM Tris, pH 7.4) containing acetyl-CoA 0.2 mM, 0.2 mM 5,5'- dithiobis-(2-nitrobenzoic acid) (DTNB), 0.55 mM oxaloacetic acid, and 0.1% Triton X-100. Final CS activity was obtained by integrating $OD^{412}$ change from 150-480 sec, and by subtracting the non-specific activity measured in the absence of oxaloacetate. COX activity was measured by detecting the decrease in absorbance at 550 nm, in a 100 mM potassium phosphate reaction buffer (pH 7.5) containing 0.1% n-dodecylmaltoside and 100 μM of purified reduced cytochrome c. Final COX activity was obtained by integrating $OD^{550}$ change over 200-600 sec and by subtracting spontaneous cyt c oxidation without cell lysate. SDH activity was measured by detecting the decrease in absorbance at 600 nm, in potassium phosphate 100 mM reaction buffer (pH 7.5) containing 2 mM EDTA, 1 mg/ml bovine serum albumin (BSA), 4 μM rotenone, 10 mM succinate, 0.25 mM potassium cyanide, 100 μM decylubuquinone, 100 μM DCIP, 200 μM ATP, 0.4 μM antimycin A. Final SDH activity was obtained by integrating $OD^{600}$ change over 200-900 sec and by subtracting activity detected in the presence of malonate (5 mM), a specific inhibitor of SDH. Complex I activity was measured by detecting the decrease in absorbance at 600 nm, in potassium phosphate 100 mM reaction buffer (pH 7.5) containing 2 mM EDTA, 3.5 mg/ml bovine serum albumin (BSA), 0.25 mM potassium cyanide, 100 μM decylubuquinone, 100 μM DCIP, 200 μM NADH, 0.4 μM antimycin A. Final Complex I activity was obtained by integrating $OD^{600}$ change over 120-600 sec and by subtracting activity detected in the presence of rotenone (500 μM) and piericidin A (200 μM), specific inhibitors of Complex I. All assays were performed at 30°C. The molar extinction coefficients used were 13.6 L $mol^{-1}cm^{-1}$ for DTNB, 29.5 L $mol^{-1}cm^{-1}$ for reduced cytochrome c, and 16.3 L $mol^{-1}cm^{-1}$ for DCIP to transform change in OD into enzyme activity.

Mitochondrial enzymatic activities were measured on a total of 340 samples, including 136 replicates of the same cell type for the same person. This provided more stable estimates of enzymatic activities than single measures would for a total of 204 individual person-cell combinations. The technical variation for each enzyme varied according to cell type, with cell types with lower enzymatic activities generally showing the highest coefficient of variation (C.V.). C.V. averaged across all cell types were: CS = 6.3%, Complex I = 16.6%, SDH = 9.3%, COX = 23.4% (Supplementary file 3).

*Mitochondrial DNA copy number*

mtDNAcn was determined as described previously (Picard et al., 2018) with minor modifications. The same homogenate used for enzymatic measurements (20 ul) was lysed in lysis buffer (100 mM Tris HCl pH 8.5, 0.5% Tween 20, and 200 ug/ml proteinase K) for 10 hours at 55°C followed by inactivation at 95°C for 10 minutes. Five ul of the lysate was directly used as template DNA for measurements of mtDNA copy number. qPCR reactions were set up in triplicates using a liquid handling station (ep-Motion5073, Eppendorf) in 384 well qPCR plates. Duplex qPCR reactions with Taqman chemistry were used to simultaneously quantify mitochondrial and nuclear amplicons in the same reactions. *Master* $Mix_1$ for ND1 (mtDNA) and B2M (nDNA) included: TaqMan Universal Master mix fast (life technologies #4444964), 300 nM of primers and 100 nM probe (ND1-Fwd: GAGCGATGGTGAGAGCTAAGGT, ND1-Rev: CCCTAAAACCCGCCACATCT, Probe:HEX-CCATCACCCTCTACATCACCGCCC-3IABkFQ.B2M-Fwd:CCAGCAGAGAATGGAAAGTCAA,B2M-Rev: TCTCTCTCCATTCTTCAGTAAGTCAACT, Probe:FAM-ATGTGTCTGGGTTTCATCCATCCGACA-3IABkFQ). *Master* $Mix_2$ for COX1 (mtDNA) and RnaseP (nDNA) included: TaqMan Universal Master Mix fast, 300 nM of primers and 100 nM probe (COX1-Fwd: CTAGCAGGTGTCTCCTCTATCT, COX1-Rev: GAGAAGTAGGACTGCTGTGATTAG, Probe: FAM-TGCCATAACCCAATACCAAACGCC-3IABkFQ. RnaseP-Fwd: AGATTTGGACCTGCGAGCG, RnaseP-Rev: GAGCGGCTGTCTCCACAAGT, Probe: FAM-TTCTGACCTGAAGGCTCTGCGCG-3IABkFQ. The samples were then cycled in a QuantStudio 7 flex qPCR instrument (Applied Biosystems Cat# 4485701) at 50°C for 2 min, 95°C for 20 sec, 95°C for 1 min, 60°C for 20 sec for 40x cycles. Reaction volumes were 20 ul. To ensure comparable Ct values across plates and assays, thresholds for fluorescence detection for both mitochondrial and nuclear amplicons were set to 0.08.

mtDNAcn was calculated using the ΔCt method. The ΔCt was obtained by subtracting the average mtDNA Ct values from the average nDNA Ct values for each pair ND1/B2M and COX1/RNaseP. Relative mitochondrial DNA copies are calculated by raising 2 to the power of the ΔCt and then multiplying by 2 to account for the diploid nature of the nuclear genome (mtDNAcn = 2ΔCt x 2). Both ND1 and COX1 yielded highly correlated mtDNAcn and the average of both amplicon pairs was used as mtDNAcn value for each sample. The overall CV across all cell subtypes was 5.1% for mtDNAcn.

*Platelet depletion in PBMCs*

For this experiment, participants were 9 community-dwelling older adults (mean age = 79, range: 64-89, 4 women and 5 men). The sample included 7 White and 2 African American participants. Exclusion criteria included diseases or disorders affecting the immune system including autoimmune diseases, cancers, immunosuppressive disorders, or chronic, severe infections; chemotherapy or radiation treatment in the 5 years prior to enrollment; unwillingness to undergo venipuncture; immunomodulatory medications including opioids and steroids; or more than two of the following classes of medications: psychotropics, anti-hypertensives, hormones replacement, or thyroid supplements. Participants were recruited from a volunteer subject pool maintained by the University of Kentucky Sanders-Brown Center on Aging. The study was conducted with the approval of the University of Kentucky Institutional Review Board. Blood (20 mL) was collected by venipuncture into heparinized tubes in the morning hours to control for potential circadian variation. PBMCs were isolated from diluted blood by density gradient centrifugation (20 min at 800 x g, brake off) using Histopaque (Sigma, St. Louis, MO). Buffy coats were washed once, and cells were counted using a hemocytometer. PBMCs (20-30M) were cryopreserved in liquid nitrogen in RPMI-1640 (Lonza) + 10% fetal bovine serum (Hyclone) + 10% DMSO (Fisher), until further processing.

For platelet depletion, PBMCs were first thawed at room temperature and centrifuged at 500 x g for 10 minutes. The supernatant was discarded, and the cells were resuspended in 2 ml of Hank's Balanced Salt Sodium (HBSS) without phenol red, calcium and magnesium (Life Technologies, Cat#14175103). Cells were then counted on the Countess II FL Automated Cell Counter (Thermo Fisher Scientific, Cat# AMQAF1000) in a 1:1 dilution with trypan blue. Cells were then divided into 2 aliquots: 1) 10 million cells for total PBMCs; 2) 11 million cells for platelet depletion. The total PBMCs were centrifuged at 2,000 x g for 2 min at 4°C and subsequently frozen as a dry pellet at -80°C until processing for enzymatic assays and qPCR. The PBMCs destined for platelet deletion cells were processed immediately.

The total PBMCs cell preparation was first immunolabeled with magnetically-coupled antibodies against the platelet marker CD61. The 11 million platelet-depleted PBMCs were then centrifuged at 300 x g for 10 minutes. After spin, the supernatant was aspirated and cells were resuspended in 80 ul of HBSS. Then 20 ul of the CD61 MicroBeads (Miltenyi Biotec, Cat# 130-051-101) were added to the cells and incubated for 15 minutes at 4°C to magnetically label platelets. Cells were washed with 2 ml of HBSS and centrifuged at 300 x g for 10 minutes. The LS column (Miltenyi Biotec, Cat# 130-042-401) was placed in the magnetic field of the MACS

Separator (Miltenyi Biotec, Cat# 130-091-051). The LS column was equilibrated with HBSS, cells resuspended in 500 ul of HBSS were applied to the LS column, and the CD61- cells were flown through the column and collected in a 15 ml collection tube. The LS column was then washed 3x with 500 ul of HBSS. The flow through was then spun at 500 x g for 10 minutes, the cell pellet was resuspended in 2 ml of HBSS and re-counted to isolate 10 M platelet-depleted cells. These cells were pelleted at 2,000 x g for 10 minutes at 4°C, the supernatant removed, and cell pellet stored at -80°C. The platelets ($CD61^+$) were recovered by flushing 1 ml of HBSS though the LS column with the plunger in a new tube, centrifuged at 3,000 x g for 10 minutes, the supernatant removed, and the cell pellet stored at -80°C until all samples could be processed for enzymatic activity assays as a single batch. For each participant, this experiment yielded three samples: 1) total PBMCs, 2) platelet depleted PBMCs, and 3) enriched platelets. Each sample was processed in parallel for RC enzymatic activity assays and mtDNAcn as described above.

*Mitotypes*

As in other complex biological systems, the function and behavior of mitochondria require synergy among multiple features. Therefore, the functional characteristics of mitochondria can be expressed by multivariate indices reflecting the inter-relations of multiple mitochondrial features. This logic is similar to body mass index (BMI), which integrates height and weight into a BMI ratio that is more easily interpretable and meaningful than either height or weight features alone. In relation to mitochondria, in human tissues the ratio of CIV and CII activities (also known as COX/SDH ratio) is used in diagnoses of mitochondrial dysfunction, whereas alone either CIV or CII activities are less easily interpretable(Picard et al., 2012). Thus, integrating primary features of mitochondrial content, genome abundance, and RC function produces integrated *mitotypes*, which reflect the functional specialization of mitochondria among different cell subtypes, or among dynamic cellular states.

Similar to BMI, the mitotypes analyzed (listed and defined in Figure 8-figure supplement 1) are computed from the ratio of two or more mitochondrial features. Each mitotype can be visualized as a scatterplot with two variables of interest as the x and y axes (Figure 8a). In the example in Figure 8a, the hypothetical cell type A has a higher value for feature 2 relative to feature 1 (Mitotype A), while cell type B has a higher value for feature 1 relative to feature 2 (Mitotype B). Note that if both features increase or decrease following the same proportions, the ratio is the same and follows a diagonal on the mitotype graph, reflecting an invariant mitotype. Thus, similar to BMI, changes in mitotypes are reflected by perpendicular movement relative to the diagonal in the mitotype space.

*Statistical analyses*

To adjust for potential order effects across the 340 samples (31 samples per 96-well plate, 17 plates total) a linear adjustment was applied to raw values enzymatic activity measures, which adjusts for potential storage and batch effects, ensuring consistency between the first and last samples assayed. Samples from both the cohort and repeat participant were processed and analyzed as a single batch.

Mann-Whitney T tests were used to compare sex differences in cell type proportions and mitochondrial measures. Throughout, effect sizes between groups were computed as Hedges' g (g) to quantify the magnitude of group differences in cell type proportions and mitochondrial measures (by sex, mitotype cell subtype and inter-individual differences). Spearman's r (r) was used to assess strength of associations between continuous variables such as age and cell proportion or age and mitochondrial measures. To assess to what extent mitochondrial features are correlated across cell subtypes (co-regulation) and to calculate the average correlation across mitotypes, Spearman's r matrixes were first computed and transformed to Fisher's Z', and then averaged before transforming back to Spearman's r ($r_{Z'}$). One-way non-parametric ANOVA Kruskal-Wallis with post-hoc Dunn's multiple comparison tests were used to compare cell type mitochondrial measures in different cell subtypes and PBMCs. Between- and within-person variation were characterized using coefficients of variation (C.V.). The root mean square of successive differences (rMSSD) was computed to quantify the magnitude of variability between successive weeks for repeated measures. Chi-square tests were computed to compare proportion of mitotype indices categories (enzyme activity per CS, enzyme ratios, enzyme per mtDNA, enzyme per mtDNA density, and enzyme per mtDNA relative to mtDNA density) by age (lower vs higher with increased age) and sex (lower vs higher in men). Finally, one-way non-parametric Friedman tests with post hoc Dunn's multiple comparisons were used to compare mitochondrial measures in platelet-depleted PBMCs, enriched platelets PBMCs, and total PBMCs. Statistical analyses were performed with Prism 8 (GraphPad, CA), R version 4.0.2 and RStudio version 1.3.1056. Statistical significance was set at $p<0.05$.
