## Appendix 2 for "Mitochondrial phenotypes in purified human immune cell subtypes and cell mixtures"

*Mitochondrial health index (MHI) among cell subtypes*

To define how RC function relates to mitochondrial content, we integrated content and function features into a single metric known as the MHI. The MHI reflects *RC capacity on a per-mitochondrion basis* (**Figure 4-figure supplement 1a**). In this study, the MHI was adapted from (Picard et al., 2018) with the addition of RC complex I (CI) activity.

Normalizing RC activities to mitochondrial content (i.e., CS) is considered the gold standard to evaluate RC dysfunction (Frazier, Vincent, Turnbull, Thorburn, & Taylor, 2020). But given the limitations of individual mitochondrial content markers for specific cell types (Larsen et al., 2012; McLaughlin et al., 2020; Picard, 2021), integrating CS and mtDNAcn into a single factor (denominator: CS+mtDNAcn) must yield a more generalizable estimate of mitochondrial content. Similarly, there may not be a single best marker of RC function across all cell types, and the RC chain operates as a multi-enzyme system that depends on all complexes. This provides the same rationale to integrate CI, CII, and CIV into a single factor (numerator: CI+CII+CIV). Integrating individually measured metrics into latent constructs also has the added advantage of triangulating measurement error, thus yielding more stable quantitative estimates of mitochondrial content and RC function. In three recent studies of chronic life stress in human PBMCs (Picard et al., 2018), ovarian tumor inflammation (Bindra et al., 2021), and mouse behavior (Rosenberg et al., 2021), we found more robust associations using MHI than individual metrics of content and RC function alone. This suggests that MHI is a functionally-relevant metric of mitochondrial biology.

The resulting MHI metric is directly interpretable. An MHI value of 100 represents the average RC activity per mitochondrion in the whole dataset. MHI values <100 reflect lower energy production capacity, whereas >100 reflect higher energy production capacity per unit of mitochondrion. For example, an individual with monocytes MHI of 80 has 20% lower RC function per unit of mitochondria than the dataset average; whereas someone with an MHI of 150 is 50% above the mean.

In our purified leukocyte dataset, MHI significantly differed between cell subtypes (ANOVA, $p<0.0001$). Monocytes had the highest MHI, which was 72% higher than B cells ($g=3.78$, $p<0.0001$), which in turn had the lowest MHI (**Figure 4-figure supplement 1b**). On average, $CD8^+$ memory T cells exhibited a 6-18% higher MHI relative to their naïve precursor ($g=1.02$, $p<0.05$), consistent with the notion that naïve and activated immune cells have different bioenergetic requirements (Nicoli et al., 2018; G. J. W. van der Windt et al., 2013). The MHI exhibited moderately consistent coherence between cell subtypes, falling in between the more coherent mitochondrial content markers (CS and mtDNAcn) and the less coherent RC markers (CI, CII, CIV) (**Figure 4-figure supplement 1c-d**).

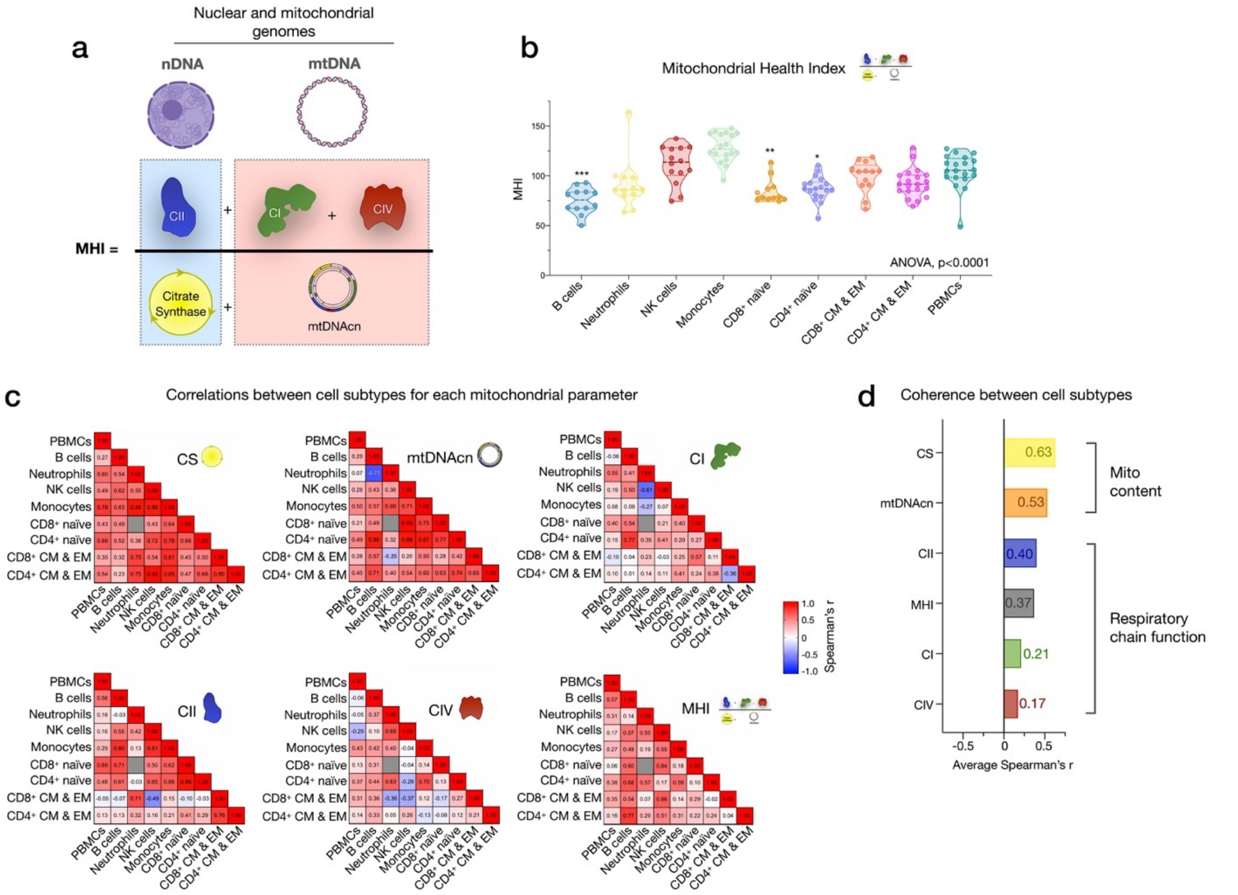


**Figure 4-figure supplement 1 – Mitochondrial health index (MHI) and coherence of mitochondrial features across cell subtypes.** (**a**) Schematic of the MHI equation reflecting respiratory chain function as the numerator, and markers of mitochondrial content as the denominator, yielding a metric of energy production capacity on a per-mitochondrion basis. (**b**) MHI across immune cell subtypes. Dashed lines are median (thick), with 25th and 75th quartiles (thin). P-values from One-Way non-parametric ANOVA Kruskal-Wallis test with Dunn's multiple comparison test of subtypes relative to PBMCs, n=12-18 per cell subtype. (**c**) Correlation matrices (Spearman's r) showing the association between cell subtypes in mitochondrial features. Correlations were not computed for cell subtype pairs with fewer than n=6 observations (gray cell). (**d**) Average effect sizes reflecting the within-person coherence of mitochondrial features across cell types (calculated using Fisher z-transformation). $p<0.05$*, $p<0.001$***.
